# Mucosal and systemic immune dynamics associated with COVID-19 outcomes: a longitudinal prospective clinical study

**DOI:** 10.1101/2023.08.04.551867

**Authors:** Mona Agrawal, Armando S. Flores-Torres, John S. Franks, Sarah Y. Lang, Thomas P. Fabrizio, Kristin E. McNair, Laura V. Boywid, Ashley J. Blair, Chloe N. Hundman, Nicholas D. Hysmith, Michael A. Whitt, Rachael Keating, Paul G. Thomas, Richard J. Webby, Amanda M. Green, Heather S. Smallwood

## Abstract

**Rationale:** COVID-19 severity varies widely; children and African Americans have low and high risk, respectively. Mechanistic data from these groups and the mucosa is lacking.

**Objectives:** To quantify mucosal and systemic viral and immune dynamics in a diverse cohort to identify mechanisms underpinning COVID-19 severity and outcome predictors.

**Methods:** In this prospective study of unvaccinated children and adults COVID-19 outcome was based on an ordinal clinical severity scale. We quantified viral RNA, antigens, antibodies, and cytokines by PCR, ELISA, and Luminex from 579 longitudinally collected blood and nasal specimens from 78 subjects including 45 women and used modeling to determine functional relationships between these data.

**Measurements and Main Results:** COVID-19 induced unique immune responses in African Americans (n=26) and children (n=20). Mild outcome was associated with more effective coordinated responses whereas moderate and severe outcomes had rapid seroconversion, significantly higher antigen, mucosal sCD40L, MCP-3, MCP-1, MIP-1α, and MIP-1β, and systemic IgA, IgM, IL-6, IL-8, IL-10, IL-15, IL-1RA, and IP-10, and uncoordinated early immune responses that went unresolved. Mucosal IL-8, IL-1β, and IFN-γ with systemic IL-1RA and IgA predicted COVID-19 outcomes.

**Conclusions:** We present novel mucosal data, biomarkers, and therapeutic targets from a diverse cohort. Based on our findings, children and African Americans with COVID-19 have significantly lower IL-6 and IL-17 levels which may reduce responsiveness to drugs targeting IL-6 and IL-17. Unregulated immune responses persisted indicating moderate to severe COVID-19 cases may require prolonged treatments. Reliance on slower acting adaptive responses may cause immune crisis for some adults who encounter a novel virus.

**At a Glance Commentary:** *Scientific Knowledge on the Subject:* Despite the disparate outcomes for African Americans and children with COVID-19 and the vital role of mucosal immunity, the majority of mechanistic clinical studies lack these groups and mucosal assessments. To date, mucosal immune responses to SARS-CoV-2 has not been adequately described and we lack data from these understudied groups.

*What This Study Adds to the Field:* This was a prospective cohort study of children and adults with confirmed COVID-19. Mortality was low (2.5%). Severity outcomes were associated with African American Race, shortness of breath, fever, respiratory disease, high blood pressure, and diabetes. We systematically characterized viral and immune factors in the mucosa and periphery and observed that moderate and severe COVID-19 were associated with longer duration, impaired clearance, early overexuberant antibody and cytokine production that was sustained. This study demonstrates that African Americans are at high risk of severe COVID-19 and display unique mucosal and peripheral immune responses. Children with COVID-19 also had distinct immune responses. This illustrates the importance of vaccination and careful clinical oversight of these populations (e.g., lower IL-6 and IL-17 levels may diminish tocilizumab, siltuximab, secukinumab, and brodalumab efficacy). This study identified generalizable outcomes predictors, systemic IL-1RA with mucosal IL-1β and IL-8, and demonstrated the utility of mucosal sampling from diverse cohorts.

## Introduction

Improving patient outcomes is hindered by the variability in COVID-19 disease manifestations. For instance, the risk of severe COVID-19 is high for African Americans (AA) and low for adolescents.(1, 2) Some studies suggested viral load induced this heterogeneity. However, viral load was lower in AA adults than Americans of European descent (EA) and was similar among pediatric and adult patients.(3, 4) Thus, nasal viral load alone does not account for the disparate outcomes in these groups. Although SARS-CoV-2 nucleocapsid (N) and spike (S) are present in the mucosa and highly antigenic,(5, 6) systemic S-specific Ab are better characterized. Peripheral S-specific IgG and IgA levels are higher in severe COVID-19.(7–9) Extensive early COVID-19 literature suggested systemic cytokine storm was linked to pathological features of COVID-19.(10, 11) Previous research in SARS-CoV-2 peripheral responses were a first step towards a more profound understanding of COVID-19. However, there is a growing consensus that there must be a mucosal immune dimension to COVID-19 that needs to be examined.(12, 13)

It is well acknowledged that the mucosa controls viral replication and invasion as well as the induction immune activation.(14, 15) Modeling indicated that adaptive responses precede the peak of mucosal infection in severe COVID-19 cases,(16) which strongly suggested mucosal immunity was involved. More evidence emerged that mucosal responses were key to controlling lung damage and mucosal interferon (IFN) levels inversely correlated with viral load and positively correlated with peripheral Ab.(17, 18) Two recent preprints indicated that viral load had no impact on the speed of recovery from COVID-19 whereas increased early mucosal immunity accelerated it; they suggested mucosal Ab were critical for timely control and clearance.(19, 20) We hypothesized that early mucosal responses to SARS-COV-2 were essential to modulating the systemic immune response and contribute to outcomes; we posit that linking the mucosal systemic responses to SARS-COV-2 could rapidly advance our mechanistic understanding as well as prognostic, therapeutic, and vaccine development. Therefore, we longitudinally collected mucosal and blood specimens from children and adults with COVID-19 to quantify SARS-CoV-2 RNA, S and N-Ag, Ag-specific IgM, IgG, and IgA, and cytokines and analyze their impact on COVID-19 outcomes to identify potential treatment targets and biomarkers.

## Methods

Additional detail on the method for making these measurements is provided in an online data supplement.

### Study Design/Participants

We conducted an IRB approved prospective observational study in Memphis TN, USA. We recruited participants from three hospitals and community COVID-19 testing sites. The study inclusion criteria required participants to have had SARS-CoV-2 testing. Non-English-speaking persons, pregnant women, and those who could not provide informed consent were excluded. Written informed consent was obtained. 78 subjects were included in this analysis. All were COVID-19 positive was based on polymerase chain reaction (PCR) or antigen tests. Cohort characteristics and treatments are summarized in Table 1.

**Table 1.** Demographics, symptoms, and clinical characteristics of the cohort by COVID-19 severity outcome. ^a^subjects with missing data. *Partial vaccination against SARS-CoV-2 (none of the participants were fully vaccinated at the time of enrollment). COVID-19: Coronavirus disease 2019; IQR: Interquartile range; AA: African Americans; EA: European Americans; U: Unknown; IVIG: Intravenous immunoglobulin. Continuous variable (age) shown as median and 75th percentiles (IQR) and one-way ANOVA test done among severity groups to find p value. Chi-square test was performed for age, sex, and racial groups (p=0·001, 0.126, and 0·0001, respectively) and outcomes with subgroups analyzed by Fisher LSD. The relationship between all other categorical variables and severity outcomes were determined with Chi-square test. Values in bold are significant (p < 0·05).

|  | Characteristics | Mild<br>n (%) | Moderate<br>n (%) | Severe<br>n (%) | Total<br>n (%) | P-value |
| --- | --- | --- | --- | --- | --- | --- |
| **Total subjects** | COVID-19 positive | 27 (34.6) | 29 (37.2) | 22 (28.2) | 78 (100) | >0.433 |
| **Age Groups** | Child | 3 (3.8) | 10 (12.8) | 7 (8.9) | 20 (25.6) | >0.432 |
|  | Adult | 23 (29.5) | 16 (20.5) | 6 (7.7) | 45 (57.7) | >0.064 |
|  | Senior | 1 (1.3) | 3 (3.8) | 9 (11.54) | 13 (16.7) | >0.371 |
| **Sex Groups** | Male | 15 (19.2) | 10 (12.8) | 14 (17.9) | 33 (42.3) | >0.574 |
|  | Female | 12 (15.4) | 19 (24.4) | 8 (10.2) | 45 (57.7) | >0.222 |
| **Racial Groups** | AA | 1 (1.3) | 9 (11.5) | 16 (20.5) | 26 (33.3) | >0.100 |
|  | EA | 10 (12.8) | 14 (17.9) | 4 (5.1) | 28 (35.9) | >0.501 |
|  | U | 16 (20.5) | 6 (7.7) | 2 (2.6) | 24 (30.8) | >0.124 |
| **Age; Median (IQR)** | Cohort | 30.4 (45) | 37 (53.5) | 50.5(63.2) | 37 (50) | 0.282 |
|  | Child | 14 (16) | 14.5 (17) | 12 (16) | 14 (16) | 0.223 |
|  | Adult | 32 (45) | 48.5 (54.2) | 45 (56.5) | 45 (50) | **<0.004** |
|  | Senior | 71 | 70 (71) | 64 (76.5) | 70 (74.5) | 0.972 |
| **Symptoms** | Rhinorrhea | 12(44.4)<sup>a=4</sup> | 7 (24)<sup>a=8</sup> | 4 (18)<sup>a=5</sup> | 23 (29)<sup>a=17</sup> | 0.159 |
|  | Anosmia | 13 (48)<sup>a=4</sup> | 7 (24)<sup>a=8</sup> | 4 (18)<sup>a=7</sup> | 24 (30.7)<sup>a=19</sup> | 0.130 |
|  | Pharyngitis | 5 (18)<sup>a=4</sup> | 4 (13.8)<sup>a=4</sup> | 1 (4.5)<sup>a=6</sup> | 10 (12.8)<sup>a=14</sup> | 0.423 |
|  | Wheezing | 1 (3.7)<sup>a=4</sup> | 4 (13.8)<sup>a=3</sup> | 4 (18)<sup>a=6</sup> | 9 (11.5)<sup>a=13</sup> | 0.177 |
|  | Shortness of breath | 2 (7.4)<sup>a=4</sup> | 15 (51.8)<sup>a=3</sup> | 17 (77)<sup>a=4</sup> | 34 (43.6)<sup>a=11</sup> | **<0.0001** |
|  | Cough | 10 (37)<sup>a=4</sup> | 16 (55.2)<sup>a=3</sup> | 14(63.6)<sup>a=4</sup> | 40 (51)<sup>a=11</sup> | 0.082 |
|  | Chills | 8 (29.6)<sup>a=4</sup> | 6 (20.7)<sup>a=4</sup> | 6 (27)<sup>a=5</sup> | 20 (25.6)<sup>a=13</sup> | 0.646 |
|  | Headache | 13 (48)<sup>a=4</sup> | 8 (27)<sup>a=4</sup> | 4 (18)<sup>a=6</sup> | 25 (32)<sup>a=14</sup> | 0.060 |
|  | Myalgia | 10 (37)<sup>a=4</sup> | 10 (34.5)<sup>a=3</sup> | 8 (36)<sup>a=4</sup> | 28 (36)<sup>a=11</sup> | 0.906 |
|  | Fever | 5 (18)<sup>a=4</sup> | 15 (51.8)<sup>a=3</sup> | 12 (54)<sup>a=4</sup> | 32 (41)<sup>a=11</sup> | **0.007** |
|  | Abdominal pain | 1 (3.7)<sup>a=4</sup> | 4 (13.8)<sup>a=3</sup> | 1 (4.5)<sup>a=4</sup> | 6 (7.7)<sup>a=11</sup> | 0.338 |
|  | Vomiting | 1 (3.7)<sup>a=4</sup> | 6 (20.7)<sup>a=3</sup> | 6 (27)<sup>a=4</sup> | 13 (16.6)<sup>a=11</sup> | 0.055 |
|  | Diarrhea | 3 (11)<sup>a=4</sup> | 7 (24)<sup>a=3</sup> | 3 (13.6)<sup>a=4</sup> | 13 (16.6)<sup>a=11</sup> | 0.455 |
|  | Chest Pain | 0 (0)<sup>a=4</sup> | 5 (17.4)<sup>a=3</sup> | 7 (31.8)<sup>a=4</sup> | 12 (15.4)<sup>a=11</sup> | **0.0006** |
| **Comorbidities** | Respiratory disease | 1 (3.7)<sup>a=3</sup> | 1 (3.4)<sup>a=8</sup> | 5 (22.7)<sup>a=10</sup> | 7 (8.9)<sup>a=21</sup> | **0.002** |
|  | Heart disease | 0 (0)<sup>a=3</sup> | 1 (3.4)<sup>a=8</sup> | 2 (9)<sup>a=10</sup> | 3 (3.8)<sup>a=21</sup> | 0.107 |
|  | High blood pressure | 0 (0)<sup>a=3</sup> | 6 (20.7)<sup>a=8</sup> | 5 (22.7)<sup>a=10</sup> | 11 (14.1)<sup>a=21</sup> | **0.0005** |
|  | Diabetes | 0 (0)<sup>a=3</sup> | 3 (10.3)<sup>a=8</sup> | 5 (22.7)<sup>a=10</sup> | 8 (10.3)<sup>a=21</sup> | **0.003** |
|  | Obesity | 2 (7.4)<sup>a=3</sup> | 5 (17.4)<sup>a=8</sup> | 5 (22.7)<sup>a=10</sup> | 12 (15.4)<sup>a=21</sup> | 0.064 |
|  | Asthma | 0 (0)<sup>a=3</sup> | 3 (10.3)<sup>a=8</sup> | 2 (9)<sup>a=10</sup> | 5 (6.4)<sup>a=21</sup> | 0.133 |
|  | Other comorbidities | 1 (3.7)<sup>a=3</sup> | 6 (20.7)<sup>a=8</sup> | 7 (31.8)<sup>a=10</sup> | 14 (17.9)<sup>a=21</sup> | **0.002** |
| **Vaccination** | Yes | 0 (0)<sup>a=11</sup> | 1*(14)<sup>a=4</sup> | 1*(4.5) | 2 (2.6)<sup>a=15</sup> | 0.700 |
| **Hospitalization** | Yes | 0 (0) | 14 (48) | 21 (95)<sup>a=1</sup> | 35 (45)<sup>a=1</sup> | **<0.0001** |
| **Therapeutics** | Steroids | 0 (0) | 8 (27) | 18 (82)<sup>a=1</sup> | 26 (33)<sup>a=1</sup> | **<0.0001** |
|  | Antiviral medication | 0 (0) | 6 (21) | 8 (36)<sup>a=2</sup> | 14 (18)<sup>a=2</sup> | **0.002** |
|  | COVID-19 Ab therapy | 0 (0) | 3 (10.3) | 5 (22.7)<sup>a=2</sup> | 8 (10.2)<sup>a=2</sup> | **0.022** |
|  | IVIg | 0 (0) | 1 (3.4) | 2 (9)<sup>a=1</sup> | 3 (3.8)<sup>a=1</sup> | 0.236 |
|  | Immune modulators | 0 (0) | 0 (0) | 1(4.5)<sup>a=2</sup> | 1 (1.28)<sup>a=2</sup> | 0.248 |
| **Survival outcome** | Survived | 27 (100)<sup>a=2</sup> | 29 (100)<sup>a=2</sup> | 20(90.9)<sup>a=1</sup> | 76 (97)<sup>a=5</sup> | 0.078 |

### Procedures

Biospecimens were obtained immediately after enrollment on visit day 1 and in follow-up visits on days 6, 14, and 28. Mid turbinate nasal swabs (MT-swabs) were collected with flocked sterile swabs inserted approximately 1 inch into the mid-turbinate region, rotated several times against the nares walls, and placed into viral transport media. Next, nasopharyngeal rinse fluids (NRF) were collected by flushing both nares with 0.1% saline, NRF was quenched with ice-cold bronchial epithelial cell growth medium and processed within 2 hours. NRF were centrifuged, cells removed, protease inhibitor cocktail added, and aliquots stored at -80°C. Blood was drawn into BD vacutainer CPT™ cell preparation tubes (BD Biosciences, Franklin Lakes, NJ, USA) with sodium citrate and processed per manufacturer guidelines the day of collection. Plasma was stored at -80°C. On study days 14 and 28, participants completed a symptom, socio-demographic, and medical history questionnaire.

### Outcomes

The outcome of COVID-19 was assigned based on retrospective chart review, completed at least 28 days from enrollment. A modified WHO COVID-19 case definition was used to assign outcome based on a standardized total severity score detailed in Table 2. Severity outcomes were classified as mild (n=27), moderate (n=29), or severe (n=22) based on cumulative points (Table 2).

**Table 2.**
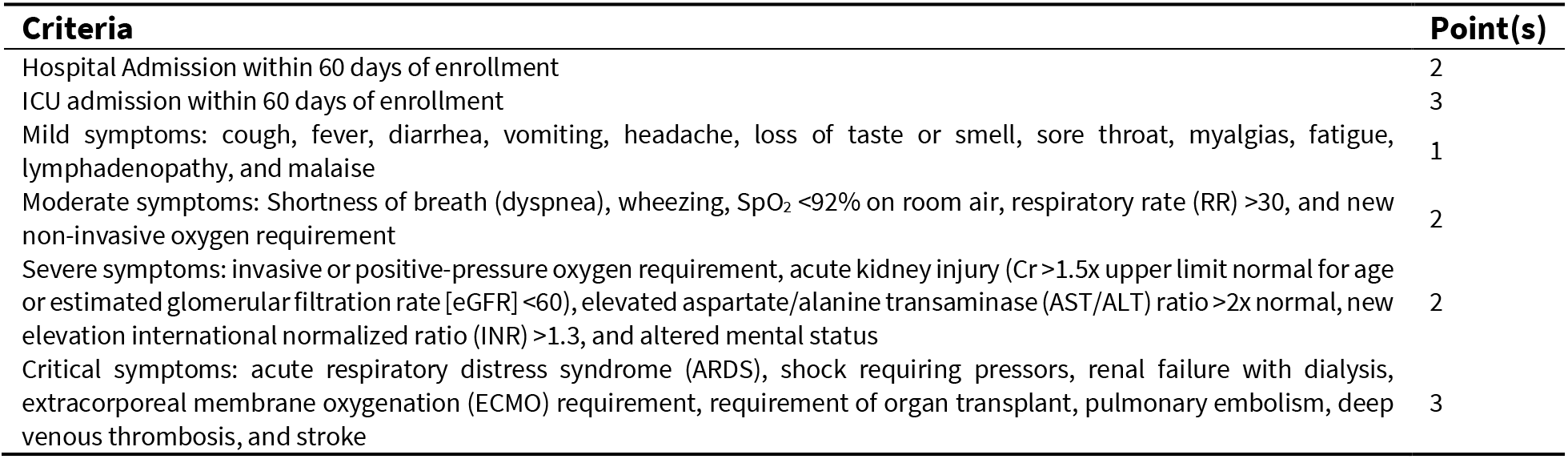
Severity Scoring Criteria. Subject severity outcomes were classified as mild, moderate, or severe COVID-19 based on cumulative point scores of ≤1, 2–4, or ≥5, respectively.

| Criteria | Point(s) |
| --- | --- |
| Hospital Admission within 60 days of enrollment | 2 |
| ICU admission within 60 days of enrollment | 3 |
| Mild symptoms: cough, fever, diarrhea, vomiting, headache, loss of taste or smell, sore throat, myalgias, fatigue, lymphadenopathy, and malaise | 1 |
| Moderate symptoms: Shortness of breath (dyspnea), wheezing, SpO<sub>2</sub> <92% on room air, respiratory rate (RR) >30, and new non-invasive oxygen requirement | 2 |
| Severe symptoms: invasive or positive-pressure oxygen requirement, acute kidney injury (Cr >1.5x upper limit normal for age or estimated glomerular filtration rate [eGFR] <60), elevated aspartate/alanine transaminase (AST/ALT) ratio >2x normal, new elevation international normalized ratio (INR) >1.3, and altered mental status | 2 |
| Critical symptoms: acute respiratory distress syndrome (ARDS), shock requiring pressors, renal failure with dialysis, extracorporeal membrane oxygenation (ECMO) requirement, requirement of organ transplant, pulmonary embolism, deep venous thrombosis, and stroke | 3 |

### Statistical Analysis

The cohort characteristics were described using the number and percent of available data for participants in the study by COVID-19 severity. Basic statistical analysis was performed using GraphPad Prism including: two-tailed Mann–Whitney U test, Log-rank Mantel-Cox test, ANOVA with Fisher’s least significant difference (LSD) procedure and posttest for linear trend, and two-way ANOVA with Tukey’s honestly significant difference (HSD) test. XLSTAT package was used for expression analysis, heatmaps, Pearson’s correlation analysis, principal component analysis (PCA), partial least squares-discriminant analysis (PLS-DA), and Classification and Regression Tree (CART) analysis. Averages with standard deviation (± SD) visualized in figures. Asterisks symbolize *P* values from these analyses as follows: *P* < 0.05 (*), *P* < 0.01 (**), *P* < 0.001 (***), and *P* < 0.0001 (****).

## Results

### Increased duration and antigen levels with higher COVID-19 severity

In this study, we analyzed 579 longitudinally collected samples from 78 COVID-19 subjects (Figures E1A and E1B). One week after COVID-19 diagnosis, no mild, 10% of moderate, and 18% of severe cases were PCR positive (Figure E2A). Mild and moderate cases significantly reduced viral load over time (Figure E2A). All subjects were infected with alpha B.1.1.7 variants (Figure E2C). N and S-Ag were quantified in NRF. Few subjects were S-Ag positive (DNS); half were N-Ag positive in week one and negative after two weeks. N-Ag was significantly higher (> 5-fold) in severe cases (Figure E2B). IgM+/IgG- and IgM+/IgG+ phenotypes are indicative of Ag exposure and development of neutralizing Ab, respectively. Most mild COVID-19 cases were IgM+/IgG-through week three and >50% were IgM+/IgG+ by week four whereas >50% of severe cases were IgM+/IgG+ in week one and all were IgM+/IgG+ after three weeks (Figure E2D). In plasma, all subjects were N-Ag negative (DNS) and severe cases had significantly higher S-Ag levels (Figure E2B). We showed that severe cases failed to reduce viral load over time and had significantly higher Ag levels, suggesting this impacted isotype switching.

### Uncontrolled mucosal and peripheral antibody responses in moderate and severe COVID-19

We quantified S and N-specific Ab responses over time (Figure E3). IgA-nasoconversion rates were faster in severe cases, but IgA levels were high early and sustained over time irrespective of severity (Figures 1A and E4A). Only mild cases increased mucosal IgG production over time (Figures 1A and E4B). We performed correlation analysis of acute phase mucosal viral load (Ct values), Ag, and Ab levels with COVID-19 outcomes. Only N-Ag and IgA had a significant relationships; positive in mild cases (*r* = 0.912, p = 0.011) and negative in severe cases (*r* = -1, *P* < 0.0001). Seroconversion rates significantly increased with COVID-19 severity (Figure 1B). Early systemic IgA, IgG, and IgM and recovery phase IgA levels also increased in proportion with severity (Figures 1B and E4C). Within the severity groups, mild significantly increased IgA and IgM and moderate increased IgM production over time (Figure E4D). Plasma microneutralization was similar irrespective of severity (Figure 1C). However, mild cases had higher neutralization potency (Figure 1C). Moreover, mild cases had strong and significant positive correlations between systemic IgG and pseudovirus and spike Ab neutralization (Figure 1D). Overall, as COVID-19 severity escalated the speed and magnitude of systemic Ab production significantly increased, which diminished neutralization.

**Figure 1.**
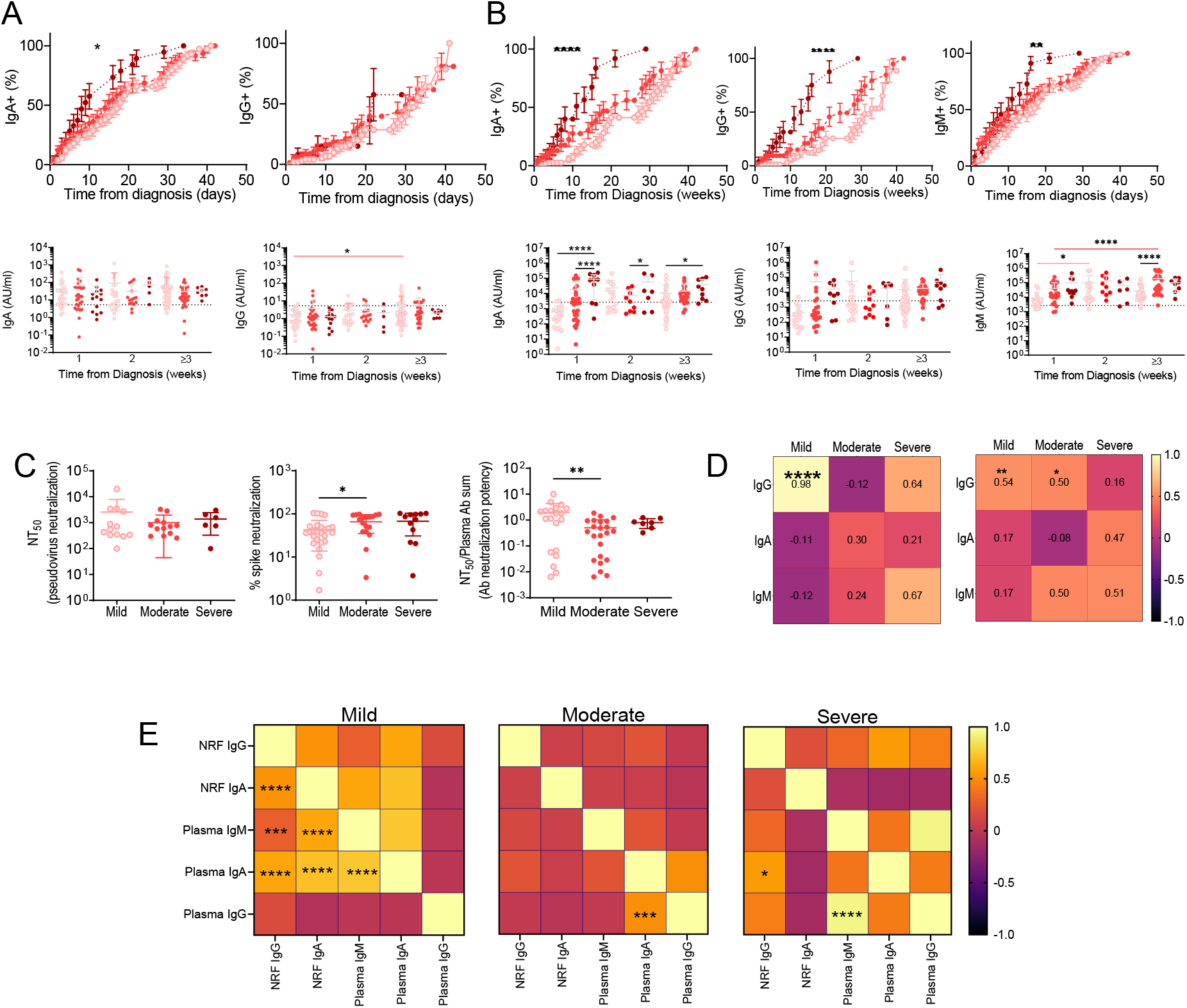
Increased rate and quantity of antibody production negatively impacts neutralization potency and coordination between local and systemic responses as COVID-19 severity escalates. N and S Ag-specific antibodies were quantitated in the NRF and plasma collected longitudinally using an ELISA. Each circle represents a patient colored by severity; mild ( ), moderate (●), and severe (●). Dotted lines represent the cut-off of positivity. Antibodies were quantified in NRF (**A**) of mild (n = 102 samples), moderate (n = 75 samples), and severe (n = 27 samples) cases. Antibodies were quantified in plasma (**B**) of mild (n = 86 samples), moderate (n = 59 samples), and severe (n = 26 samples) cases.Nasoconversion (**A**) and seroconversion (**B**) rates were compared by Log-rank Mantel-Cox test. Antibody magnitudes in the severity groups by weeks following diagnosis were compared by one-way ANOVA with Tukey’s HSD test. **C** 50% pseudovirus neutralization (NT_50_) neutralization was determined in plasma samples collected at study day 28 (mild [n = 13], moderate [n = 13], and severe [n = 6]). Percent spike Ab neutralization determined in plasma samples collected plasma between 14 and 42 days from after diagnosis (mild [n = 26], moderate [n = 18] and severe [n = 12]). Ab neutralization potency (NT50/sum of Ag-specific Ab) in plasma samples from mild (n = 20), moderate (n = 23), and severe (n = 7) outcome groups. Severity groups were compared by one-way ANOVA with Tukey’s HSD test. **D** Correlation matrix for plasma antibody levels and NT_50_ (left) or spike neutralization (right) are depicted with Pearson’s coefficient inset. **E** Correlation matrix for Ag-specific antibody magnitudes in plasma and NRF is depicted for mild (left), moderate (middle) and severe (right) outcome groups. 130 paired NRF and plasma samples were analyzed by two tailed Pearson’s correlation test. Correlation index scale depicted on the right. **A-E** Significance levels are indicated by asterisks (\**P* < 0.05, \*\**P* ≤ 0.01 and \*\*\*\**P* ≤ 0.0001).

Only the mild group had well-coordinated mucosal and systemic Ab responses, including significant and strong relationships between IgA levels in NRF and plasma (*r* > 0.7, p<0.0001) and plasma IgM with mucosal IgA and IgG (*r* > 0.6, p<0.0001) (Figures 1E and E5A). Expression analysis with hierarchical clustering was performed on paired NRF and plasma samples (Figures E5B,C and Table E1). Most mild cases displayed similar coordinated Ab responses in the first two weeks as well as in recovery (Figures E5B cluster 1 and E5C cluster 1c). However, seven mild cases had uniquely high Ab profiles within two weeks and five at week three or more; three cases carried over (Figures E5B and E5C; clusters 2, 3, 4, and 5 and clusters 1a, 1b, and 2, respectively). One-fourth of moderate sample pairs had high mucosal and/or systemic Ab (clusters 1b and 2), the remaining subjects had lower systemic Ab levels (cluster 1a). Severe cases clustered by high and low systemic IgA (Figure E5B). Intubation prohibited NRF collection limiting paired samples from severe cases. Nonetheless, at these later times, paired samples from moderate or severe cases tended to have off or high global Ab responses or displayed heterogeneous mixes within individuals of high and low Ab that were uncorrelated (Figures E5C and Table E1). Overall, mild COVID-19 was associated with slower well-coordinated mucosal and systemic Ab responses whereas moderate and severe outcomes were associated with premature simultaneous Ab overproduction with a loss of control between compartments and evidence of uncorrelated responses within individuals.

### Higher inflammatory mucosal microenvironment and uncoupled systemic responses associated with increased COVID-19 severity

Cytokines were quantified in NRF and plasma (Figures E6A and E7A). Sankey diagrams are used in physics to represent energy transfers between processes where inputs can be traced through a series of events or stages to outputs. We employed this technique to illustrate the amount of growth factor, chemokine, and cytokine inputs traced through the magnitude of each immune factor and their convergence to their main cellular target through to adaptive, innate, and non-immune response stages (Figures 2 and 3). As COVID-19 severity increased so did the levels of mucosal cytokines targeting non-immune cells while the concentration of chemokines that target dendritic, natural killer, and T cells decreased (Figure 2). There were very strong and significant positive correlations (*r* > 0.8, *P* < 0.0001) between mucosal cytokines in mild, moderate, and severe outcome groups (Figure E6B). Of these, several had significant and major differences between the mild and severe outcome groups (Figure E6C). Differential expression analysis showed as COVID-19 severity increased IL-8, MDC, MCP-1, MCP-3, GM-CSF, Flt-3L, IL-1β, TNF-β, IL-12p40, and IL-2 significantly increased (Figures E6D-G). After this acute phase, MCP-3, MIP-1α, MIP-1β, RANTES, GM-CSF, Flt-3L, PDGF, G-CSF, IL-2, IL-6, IL-12p70, sCD40L, TNF-α, and IL-10 significantly increased (Figures E6D-G). IP-10 and IFN-α2 significantly decreased progressively as severity increased in the acute phase (Figure E6D and E6F).

**Figure 2.**
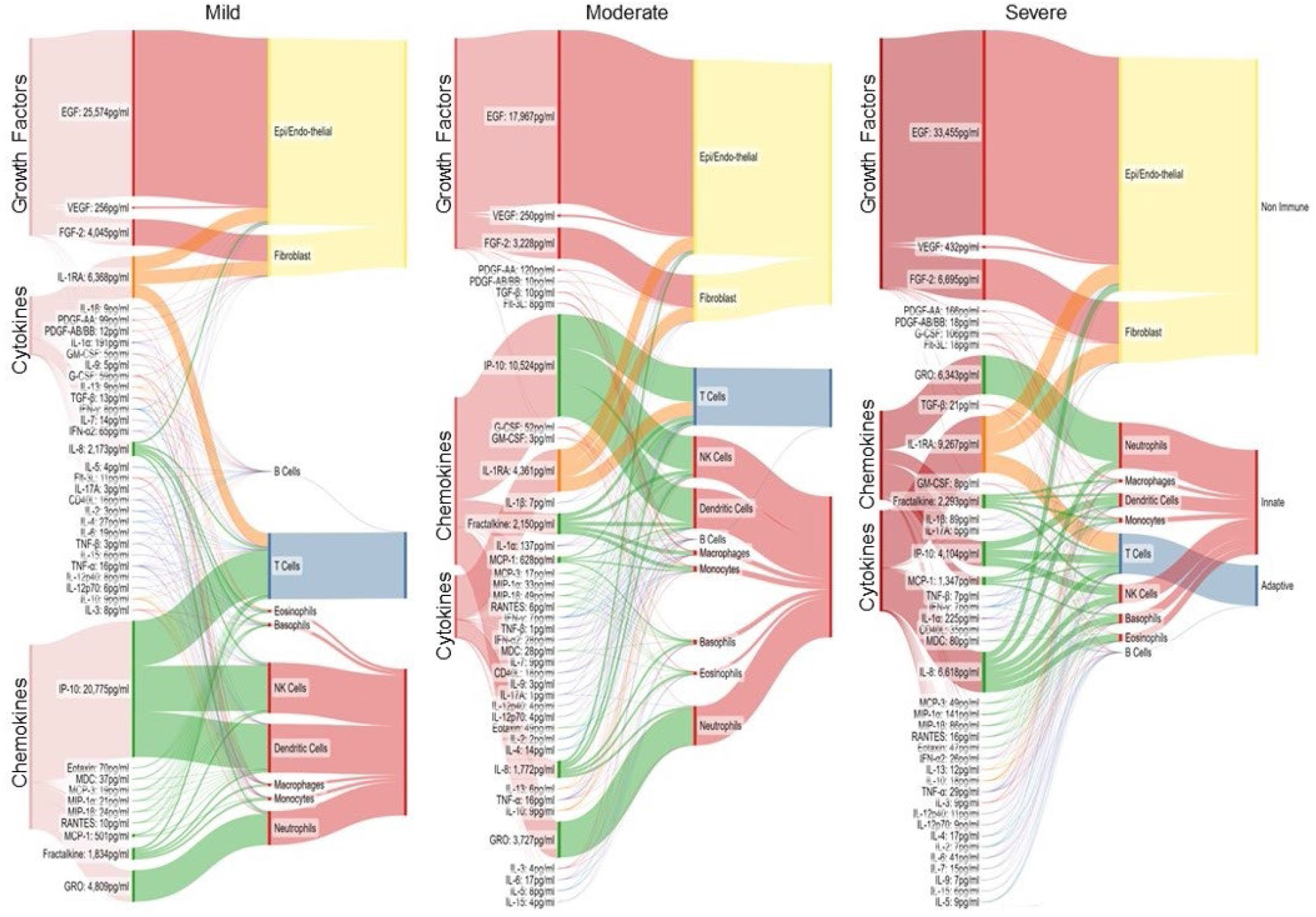
Early mucosal growth factors, chemokines, and cytokines targeting dendritic, NK, and T cells are diminished as COVID-19 severity increases. Immune factors were quantified in NRF from 61 subjects (mild n = 26, moderate n = 25, and severe n = 10) and by Luminex. Sankey diagrams depict the magnitude of cytokines, growth factors, and chemokines flowing to cell targets in each severity group during the acute phase of COVID-19 (≤7 days from diagnosis). The mean pg/ml of each immune factor was input into the model to generate chord thickness. Chords were colored by function: growth factors (red), cytokines (orange), and chemokines (green), anti-inflammatory (orange), adaptive immunity (blue) and pro-inflammatory (purple). Immune factors were linked to their major cellular targets in the context of viral infection. Cell targets were grouped into non-immune (yellow; epithelial, endothelial and fibroblasts), innate (red; eosinophils, basophils, neutrophils, NK-cells, dendritic cells, macrophages, and monocytes), and adaptive (blue; T and B-cells). The band width per cell type corresponds to the total of cytokines directed at each cell target.

As a part of a larger study, that combined two clinical cohorts, we quantified cytokines in plasma of healthy and SARS-CoV-2 infected.(21) For facile comparison with mucosal cytokines, we extracted the data from subjects in this study (Figure E7A). During the acute phase, as COVID-19 severity increased the levels of systemic cytokines targeting dendritic, NK, and T-cells increased while those targeting neutrophils, basophils, and eosinophils decreased (Figure 3). There were very strong significant positive correlations observed between systemic cytokines (*r* > 0.8, *P* < .0001). Levels of 13 cytokines were significantly different among COVID-19 severity outcomes irrespective of duration (Figure E7B). Of these, IL-1RA, IL-4, IL-6, and MCP-1 had the largest increased fold change while IL-1α significantly decreased (Figure E7C). Moderate and severe groups had significantly higher levels of cytokines compared to mild cases, including IL-1RA, IL-8, IL-6, IL-10, IP-10, IL-15, and MIP-1β within one week of diagnosis after which IL-1RA, IL-8, IL-15, and MIP-1β remained elevated (Figure E7D).

**Figure 3.**
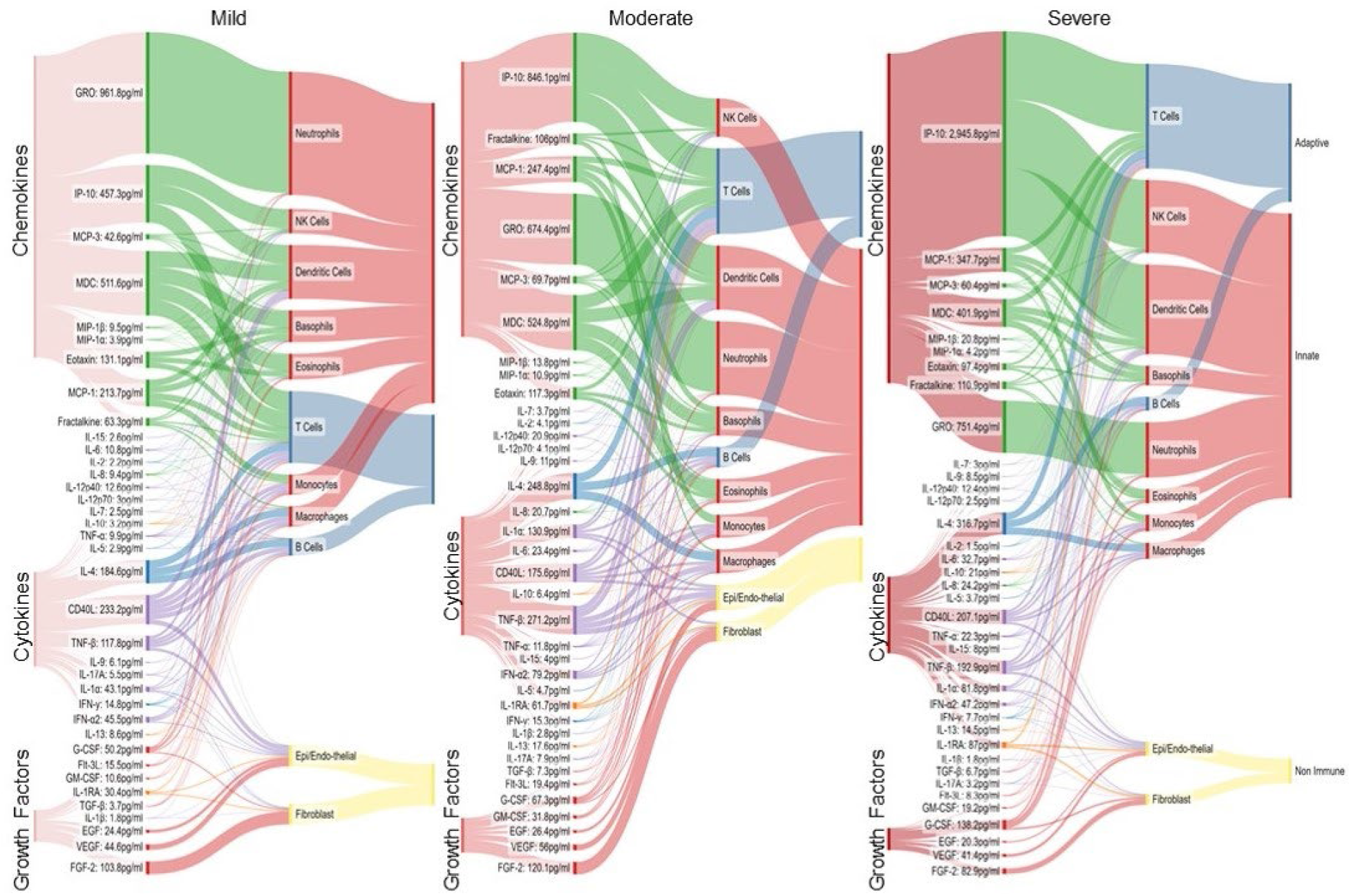
Systemic immune factors targeting neutrophils decreased while those targeting dendritic, NK, and T cells increased as COVID-19 severity increased. Immune factors were quantified in plasma from 59 subjects (mild n = 26, moderate n = 22 and severe n = 11) by Luminex. Sankey diagrams depict the magnitude of cytokines, growth factors, and chemokines flowing to cell targets in each severity group during the acute phase of COVID-19 (≤7 days from diagnosis). The mean pg/ml of each immune factor was input into the model to generate chord thickness. Chords were colored by function: growth factors (red), cytokines (orange), and chemokines (green), anti-inflammatory (orange), adaptive immunity (blue) and pro-inflammatory (purple). Immune factors were linked to their major cellular targets in the context of viral infection. Cell targets were grouped into non-immune (yellow; epithelial, endothelial and fibroblasts), innate (red; eosinophils, basophils, neutrophils, NK-cells, dendritic cells, macrophages, and monocytes), and adaptive (blue; T and B-cells). The band width per cell type corresponds to the total of cytokines directed at each cell target.

IL-6, IL-8, IL-10, and MIP-1β significantly increased with severity in NRF and plasma (Figures E6 and E7). NRF had higher levels of proinflammatory cytokines that initiate proliferation and infiltration, such as EGF, FGF-2, VEGF, G-CSF, IL-8, IP-10, GRO, and MIP-1α, whereas plasma had higher levels of adaptive and proinflammatory cytokines, such as IL-4, CD40L, TNF-β, and IFN-α2 (Figures E6 and E7). In contrast to moderate and severe outcomes, mucosal and systemic cytokine levels in mild cases correlated well displaying positive relationships in the acute phase which dissipated in the recovery phase (Figure 4A). Correlation coefficients of cytokines with significant relationships between NRF and plasma were plotted (Figures 4B-D). In the acute phase, mild cases had far more significant positive associations between compartments with notably more mucosal growth factors and adaptive cytokines that correlated to systemic

**Figure 4.**
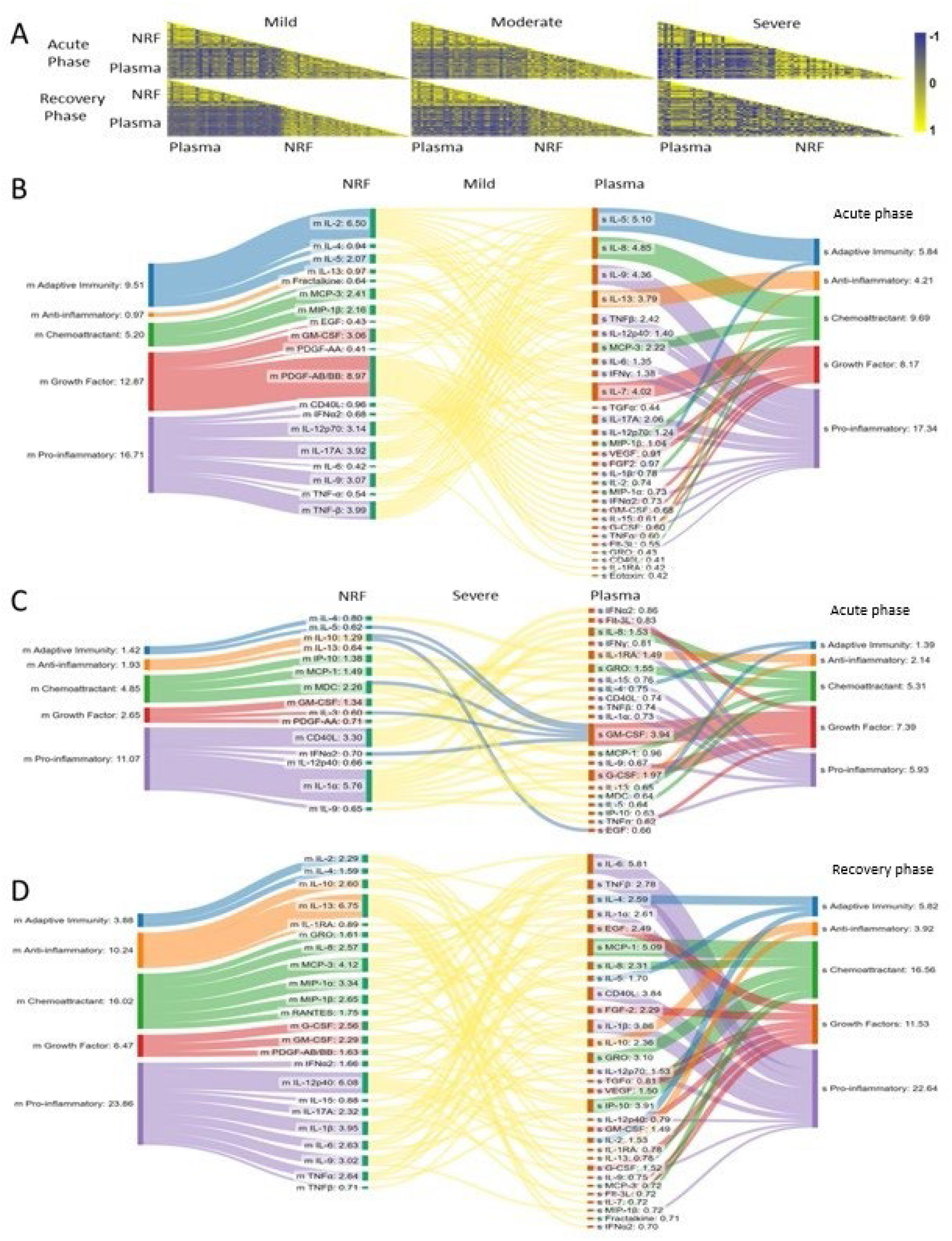
Correlation between mucosal and systemic cytokine responses by COVID-19 severity outcomes. The linear correlation between mucosal and systemic cytokines within each severity group was measured in the acute and recovery phases (≤7 days and ≥8 days from diagnosis, respectively). **A** Heat maps were colored by Pearson correlation coefficients. Positive correlations depicted in yellow and negative in blue and correlation index scale is depicted at the right of correlation plots ranging from -1 to 1. **B-D** Each cytokine with significant correlations (*p* < 0.05) and absolute value of *r* > 0.4 between NRF and plasma was mapped between compartments. The absolute value of Pearson correlation coefficients were input into the model. Nodes for mucosal (m) and systemic (s) compartments were colored green and orange, respectively, and were connected by chords colored in yellow for positive and blue for negative correlations. Chord thickness depicts the total *r* value. The cytokines were linked to their major functional groups: growth factors (red), cytokines (orange), and chemokines (green), anti-inflammatory (orange), adaptive immunity (blue) and pro-inflammatory (purple). Cumulative *r* values for each functional group corresponds to the starting and end node thickness. The flow of cytokines between mucosal and systemic compartments were depicted for acute phase mild (**B**), acute phase severe (**C**), and recovery phase severe (**D**). Mild cases lacked significant correlations with absolute value of r > 0.4 in the recovery phase.

cytokine levels compared to severe cases, (Figures 4B and 4C). Severe cases also had six mucosal cytokines that negatively correlated with systemic GM-CSF and EGF (Figure 4C). After the acute phase, mild cases lacked significant correlations between mucosal and systemic cytokines (*r* < 0.4). In contrast, the number of significant positive correlations between compartments in the severe group increased going from the acute to recovery phases (Figure 4D). Our results demonstrated that moderate and severe COVID-19 outcomes were associated with increased cytokine production and altered cytokine profiles and cellular targets that promoted mucosal and systemic proinflammatory states and loss of temporal control, which suggested a failure to resolve or restrain the immune response.

### Unique immune responses to COVID-19 and outcome predictors

Severe outcome had no correlation with sex and very weak correlation with age (*r* = 0.187). Subjects were grouped into child, adult, and senior by age (≤ 19, 20-60, and ≥61 years, respectively). When compared to adults, children with COVID-19 had significantly more IL-8 (>18-fold) and >2-fold more IL-4, IL-6, IL-15, and TNF-β and less IL-10 in plasma (Figure 5A). Within moderate and severe outcomes, children had significantly higher systemic S-Ag, IgA, CD40L, fractalkine, G-CSF, IL-1RA, MCP-3, IL-4, and mucosal IL-1β, and MDC with lower spike neutralization titers and mucosal IFN-α2 and fractalkine compared to adults (Table E2). Seniors also had significant differences in systemic and mucosal immune factors compared to adults (Figure 5A). AA had significantly lower age and systemic IFN-α2 and MIP-1β with higher IgA, IL-15, IP-10, IL-10, IL-2, TGF-α, and MIP-1α compared to their EA counterparts (Figure 5A). In the mucosa, IL-1α, MCP-1, MCP-3, PDGF-AB/BB, and FGF-2 were significantly higher while IL-17A and IL-2 were around 46 and 5-fold lower in AA than in EA (Figure 5A). Overall, children, seniors, and African Americans with COVID-19 had distinct immune responses to SARS-CoV-2.

**Figure 5.**
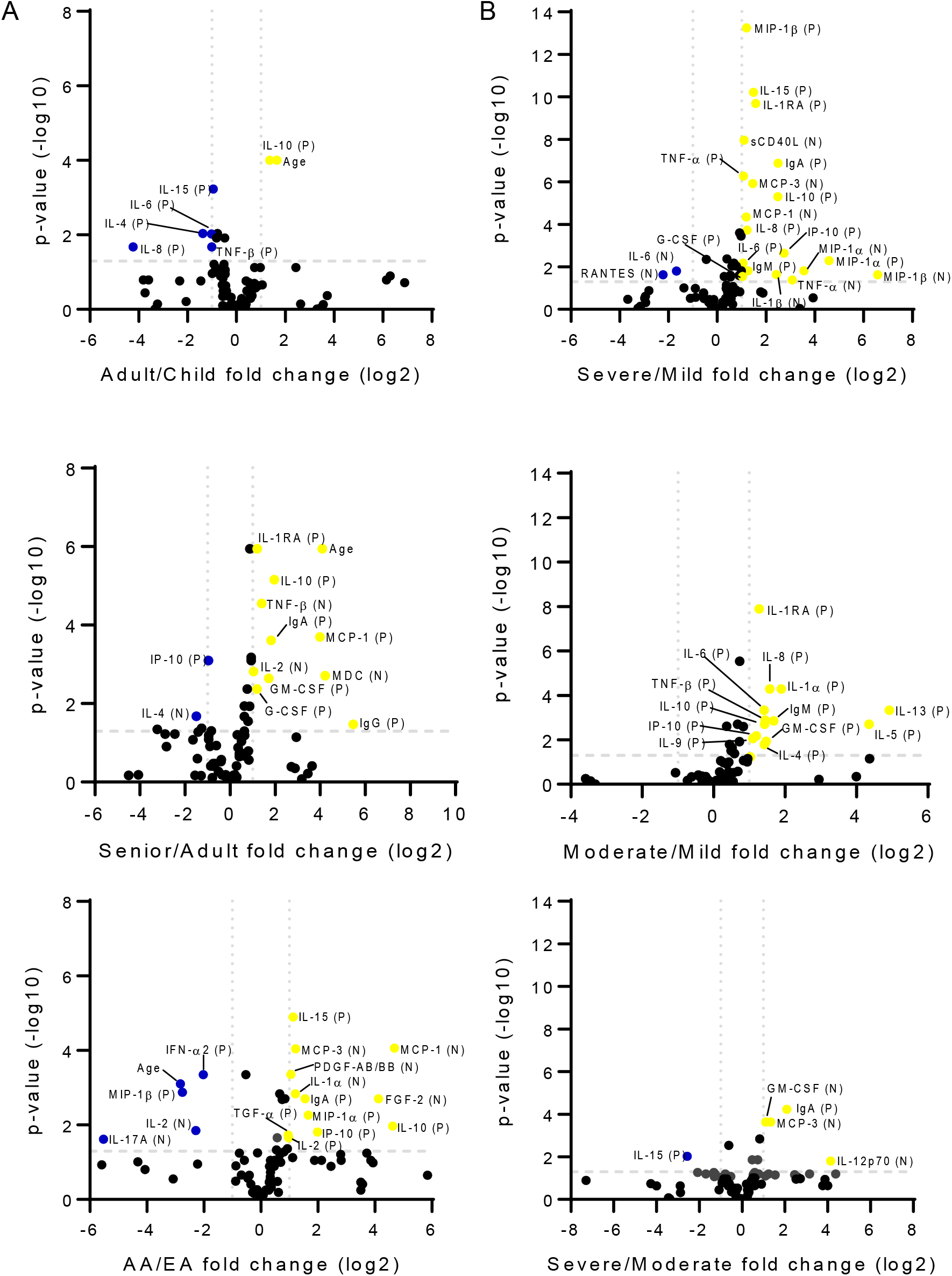
Host, viral, and immune factors associated with age, race, and severity. **A** Correlates identified by age group (left and middle) or ethnicity (right). **B** Correlates identified based on COVID-19 disease severity. Total eighty four biomarkers together in NRF and plasma were quantified and analyzed for differential expression between two groups as indicated in the plots; Adult (n = 159) vs Child (n = 45); Senior (n = 30) vs Adult (n = 159); AA (n = 60) vs EA (n = 87); Moderate (n = 80) vs Mild (n = 103); Severe (n = 51) vs Mild (n = 103) and Severe (n = 51) vs Moderate (n = 80) using XLSTAT. Differentially expressed biomarkers were plotted as volcano plot using GraphPad software. Out of 84 biomarkers, around 30 biomarkers differentially expressed between two groups with lowest *P* values. Y-axis represents the -log10 *P* values and X-axis log2 fold change of biomarkers between two groups. Dashed lines at y and x-axis showed the cut-off for significant fold increase or decrease. Each dot represents a biomarker. Factors that significantly increased are colored yellow, significantly decreased blue and no significant change with black color. Biomarker differentially expressed in NRF and plasma are symbolized with N and P respectively, right to the biomarker name in the brackets.

The entire cohort was used to identify biomarkers and predictors of severity outcomes. Severe COVID-19 cases had significantly higher sCD40L, MCP-3, MCP-1, MIP-1α, MIP-1β, IL-1β, and TNF-α in the mucosa and MIP-1β, IL-15, IL-1RA, TNF-α, IL-10, IL-8, IP-10, IL-6, MIP-1α, G-CSF, IgA, and IgM in plasma compared to mild cases (Figure 5B). However, mucosal IL-6 and RANTES were significantly higher in the mild group compared to severe. Moderate cases also had significantly higher systemic cytokines and IgM compared to mild cases (Figure 5B). PLS-DA analysis was performed on all viral and host quantitative and qualitative explanatory variables (Figure E8). Severe COVID-19 correlated with concentrations of mucosal MCP-3, GM-CSF, TGF-α, IL-1β, IL-2, IL-8, IL-9, and IL-17A along with systemic MIP-1β, IP-10, IL-10, and IgA as well as AA ethnicity, age, nasal S Ag positivity, IgG positivity and IgM+/IgG+ phenotype (Figure E8A). The mild and severe groups were well discriminated with some overlap in the moderate group (Figure E8B). Supervised machine learning algorithms were used to explain and predict outcomes on the basis of viral load, Ag, Ab, and cytokine levels. Briefly, we found if systemic IL-1RA was low then severity was mild in 71 out of 103 cases. When IL-1RA levels were high, severity was moderate in 75 out of 80 and severe in 46 out of 50 cases, and when accompanied by high mucosal IL-8 all cases were severe (Figure E9A and Table E3). When restricted to week one observations, mucosal IL-1β, IFN-γ, and CD40 along with systemic IL-1RA and IgA had predictive value (Figure E9B and Table E4). From these results, it is clear that mucosal assessments added value to the identification of severity biomarkers and outcome predictors.

## Discussion

We investigated whether disparate outcomes for COVID-19 can be partly explained by mucosal immunity and feedback between local and peripheral responses. We examined this in a cohort that included understudied populations. This prospective longitudinal study adopted a comprehensive approach to quantify severity outcomes, virus, and immune responses in mucosal and blood samples. A challenging analytical problem which arises in this domain are confounding variables. To address this, we studied naïve unvaccinated subjects with alpha strain infections that were enrolled within 72 hours of diagnosis. We verified 80% were in the same stage of infection based on IgM seropositivity in week one. This IgM positivity rate agreed with meta-analysis of 52 cohorts, which established this was remarkably accurate for evaluating infection phase.(22) We identified unique responses in children, seniors, and AA with COVID-19; defects in mucosal and systemic immune responses associated with severity; and sets of severity biomarkers and outcome predictors. It is notable these results came from a representative cohort of children and adults with a wide range of COVID-19 severity that included equal representation of AA and females. They were recruited to reduce bias, facilitate stratified analysis of subgroups, and increase the generalizability of this study. To our knowledge, this is the first mucosal immunity assessment in such a diverse cohort and first reported comparison of mucosal and peripheral immune kinetics extending over a month.

We showed that acute phase viral load and mucosal IgA production were similar irrespective of outcome. We describe lingering virus and significantly higher mucosal N-Ag in moderate and severe cases compared to mild cases, which suggests both mucosal IgA and IgG are needed for clearance. We observed systemic S-Ag levels, the speed of class switching, seroconversion rates, and early Ag specific Ab levels significantly increased proportionally with severity outcomes. These results go beyond previous reports, showing seroconversion increased with severity.(23, 24) Our results showed that mild cases slowly increased S and N-Ag specific Ab production over time improving their neutralization potency. A similar conclusion was reached by Hendriks et al. when they showed Ab affinity decreased as COVID-19 severity increased.(25) Another promising finding in mild cases was that both spike and micro-neutralization significantly positively correlated with systemic IgG levels, which suggest IgG was neutralizing. Our results cast new light on early vigorous systemic Ab responses in moderate and severe COVID-19 cases that are less effective and are driven by Ag and/or cytokines, as opposed to viral load.

Our cytokine results demonstrated that the magnitude of local and systemic responses increased proportionally with severity outcomes. Mucosal IL-1β, IL-2, IL-8, GM-CSF, MCP-1, MCP-3, MDC, and TNF-β rapidly increased and cytokine profiles shifted to proliferative growth factors as severity escalated. A similar pattern of results was obtained in bronchoalveolar lavages.(26, 27) A novel finding was that mucosal and systemic cytokines were only coregulated in the acute phase for mild cases whereas severe cases had an increase in cytokine correlations after week one, which included many proinflammatory factors. Overall, our analysis found evidence of an early transition to a highly inflammatory mucosal microenvironment with sustained positive feedback between local and systemic proinflammatory responses that was associated with poor COVID-19 outcomes. Our observations are consistent with the huge influx of activated immune cells into the respiratory tract and immunopathologies associated with severe COVID-19. At this stage of understanding, we believe increasing catabolism of pro-inflammatory cytokines, quenching Ag and activated proinflammatory cells, and discrete targeting of mucosal and systemic responses may be beneficial. Moreover, our data indicated this state was protracted and suggests extending treatments that manipulate it.

Planned comparisons revealed unique immune responses in children and AA with COVID-19. We observed significantly more systemic IL-4, IL-6, IL-8, IL-15, and TNF-β and less IL-10 in children compared to adults with COVID-19. It is notable that children had 18-fold higher IL-8 and 2.6-fold lower IL-10 compared to adults whereas IL-8 and IL-10 were 2.3 and 5.3-fold higher, respectively, in severe compared to mild cases overall. Apart from IL-10, these cytokines mainly induce infiltration, proliferation, and activation of B-cells, T cells, NK-cells, neutrophils, and phagocytes, which shows evidence of robust innate responses in children. In contrast, AA had significantly higher severity scores (mean difference = 3.453, *P* < 0.0001); significantly less systemic IFN-α2 and MIP-1β (-4 and -6.7 -fold, respectively); and significantly lower mucosal IL-17A and IL-2 and higher mucosal IL-1α, PDGF-AB/BB and FGF-2 (-46, -4.9, 2.3, 2 and 17.4 -fold, respectively). Importantly, systemic IFN-α2 and MIP-1β were the opposite (increased by 2 and 2.3-fold, respectively) and mucosal cytokines absent in severe compared to mild cases overall. Our findings of lower systemic IFN-α2 and MIP-1β and mucosal IL-17A and IL-2 in AA at least hint that anti-viral signaling and macrophage, DC, T cell, and NK cell chemotaxis are weakened in the periphery with diminished monocyte and neutrophil recruitment and helper T cell responses at the site of infection. Together our data indicate children and AA have specific immune responses to SARS-CoV-2, and suggest existing biomarkers derived from EA adults may not be as accurate for these patient groups.

When combined, our findings support several non-mutually exclusive potential contributing factors to COVID-19 outcomes. IgA may be detrimental or beneficial. Conflicting reports suggested mucosal S-specific IgA correlated with mortality while others showed it correlated with mild COVID-19, delayed or absent systemic IgA, and inversely correlated with age. (15, 28) In the mucosa, we observed IgA was elevated early irrespective of severity while IgG was increasing in mild cases, which suggests IgA alone does not account for desperate outcomes. High systemic IgA levels blocked IgG Ag-binding, neutralization, and protective functions after HIV vaccination.(29) In line with previous findings,(25) we observed moderate and severe COVID-19 cases produced significantly more systemic IgA. This was very early; IgA was undetectable in mild cases before week 2. Thus, IgA interference may occur in moderate and severe COVID-19 and may reduce their neutralization capacity, but given mild cases lacked early mucosal and systemic IgA and IgG, any impact on outcome would likely come later.

Alternatively, individuals who mount more specific initial responses to SARS-CoV-2 may develop milder disease. This notion is also supported by our observation that mild cases showed higher neutralization potency. However, neutralizing Ab alone are insufficient for viral clearance so this would not explain the uniquely mild disease presentation in children. Finally, a quality-quantity trade-off may occur early in the response to SARS-CoV-2, such that a rapid and robust humoral, cytokine, and cellular response is initiated at the expense of a slower and more specific response. This is supported by our observations that the mucosal and systemic Ag loads, Ab levels, and proinflammatory cytokine levels rapidly accumulate to significantly higher levels in moderate and severe cases. High Ag levels drive production of proinflammatory cytokines and cells, all of which converge to support fast-acting extrafollicular synthesis of lower affinity and more broadly reactive Ab, stemming in part from more random naïve B-Cell selection. This is consistent with our observation of very high levels of less potent Ab and other reports of COVID-19 inducing Ab against IFN.(30) Moreover, adults rely on highly specific adaptive immune responses whereas children frequently encounter new viruses and generally depend on innate immunity. Directly in line with previous findings,(31) we demonstrated that children had stronger innate responses to SARS-CoV-2 than adults, which we also showed was true among just moderate and severe cases. This suggests that adults may be at a disadvantage when encountering a novel virus. From this standpoint, at the onset of COVID-19 some adults may enter an immune crisis whereby fast and strong immune responses are selected at the expense of precision, which furthers collateral damage and inflammation that in turn reinforce signals that make this crisis harder to resolve.

Based on observations in this cohort we conclude mucosal responses are protective and major contributing factors to COVID-19 severity. We reveal new findings that in moderate and severe cases have extremely rapid and robust humoral responses leading to less effective neutralization driven by local and peripheral Ag and cytokine levels. Our comparative analysis over time revealed the importance of early coordination between mucosal and systemic responses, which are distinct yet interdependent and require well-regulated timing and well-balanced magnitudes to prevent immunopathologies associated with poor outcomes; this coordination may be an essential determining factor in COVID-19 disease progression. Collectively, these data provide regulatory nodes that can be exploited for therapeutic purposes and mechanistic insights into mucosal responses to SARS-CoV-2. Our study suggests that improving mucosal responses should be a priority and supports therapeutic targeting of the mucosa and including mucosal fluids when identifying biomarkers, correlates of protection, and outcome predictors for COVID-19. Our results provide a basis for a set of severity biomarkers for use in both mucosal and blood samples (IL-2, IL-6, IL-8, IL-1RA, MIP-1β, and GM-CSF) and the use of mucosal IL-8, IL-1β, and IFN-γ with systemic IL-1RA and IgA for tracking and predicting outcomes over the course of COVID-19. We acknowledge larger cohorts with other minorities are needed. Nonetheless, these data may help improve outcomes for understudied groups. Future research could examine the cause of abnormal immune kinetics in moderate and severe cases, IgA interference, extrafollicular of Ab production, mucosal cell specific responses as well as drivers of immune responses subpopulations with unique COVID-19 outcomes such as children and AA.

## Acknowledgements

We are grateful to the subjects, their families, and our collaborating hospitals and their staff whose cooperation made this study possible. We would also like to recognize the effort of the members of the UTHSC Institutional Review Board who reviewed, improved, and approved the human participant research required for this study. Plasmid expressing codon-optimized cDNA for the SARS-CoV-2 spike was generously provided to us by Dr. Florian Krammer, Icahn School of Medicine, Mount Sinai. Additionally, we would like to thank Andrew Gienapp for editing this manuscript, Manasa Mallampaty for assistance in integrating our consents and questionnaires into RedCap, and Kerry Moore and Dr. Sandra Arnold for clinical study feedback and compliance oversight.

## Funding

This project has been funded in part with Federal funds from the National Institute of Allergy and Infectious Diseases, National Institutes of Health, Department of Health and Human Services, under Contract No. 75N93021C00016. Other funding sources include Le Bonheur Children’s Hospital Research Grants, the UTHSC Office of Research, and the Children’s Foundation Research Institute.

**Footnotes**

## Notes

### Competing Interest Statement

The authors have declared no competing interest.

