## Supplement data for "Mucosal and systemic immune dynamics associated with COVID-19 outcomes: a longitudinal prospective clinical study"

**ONLINE DATA SUPPLEMENT**

**Supplementary Methods**

**Regulatory compliance:** Written consent was provided from participants or their legal guardians in compliance with 45 CFR46 and the Declaration of Helsinki regarding ethical principles for medical research. Electronic data from medical records were extracted as per Redcap data guidelines without patient identity.

**Inclusion and exclusion criteria:** Non-English-speaking persons, pregnant women, and those who could not provide informed consent or lacked an available designated decision maker were excluded from the study. Participants visited to local hospitals or to the UTHSC’s community COVID-19 testing site to have had SARS-CoV-2 testing due to symptoms or known exposure to COVID-19 positive individuals were included in the study.

**Human Subjects and Specimens:** The Institutional Review Boards of The University of Tennessee Health Science Center (UTHSC) and St. Jude Children’s Research Hospital approved this study. Participants or their legal guardians provided written informed consent in compliance with 45 CFR46 and the Declaration of Helsinki regarding ethical principles for medical research. Participants were recruited from Le Bonheur Children’s Hospital, Methodist Germantown Hospital, Methodist University Hospital, and UTHSC’s outpatient community COVID-19 testing sites. The study inclusion criteria required participants to have had SARS-CoV-2 testing due to symptoms or known exposure to COVID-19 positive individuals. Participants were considered COVID-19 positive if SARS-CoV-2 polymerase chain reaction (PCR) tests conducted in clinical laboratory was positive. Non-English-speaking persons, pregnant women, and those who could not provide informed consent or lacked an available designated decision maker were excluded from the study.

Demographic, epidemiological, clinical, laboratory, treatment, and outcome data were collected from electronic medical records as available. Clinical features were collected using a standardized data record form. The Severity of COVID-19 illness was assigned based on retrospective chart review, completed within 28 days from enrollment. A modified World Health Organization (WHO) COVID-19 case definition was used to assign severities of asymptomatic, mild, moderate, and severe. Standardized point values were assigned based on hospital admission, intensive care unit (ICU) admission, and symptoms detailed in (**Table 2**). Participants were classified with mild, moderate, or severe COVID-19 based on cumulative point scores (i.e., total ≤1, 2–4, or ≥5, respectively). Hospitalized participants were all designated as moderate or severe, and all participants admitted to the ICU were designated as severe.

Specimens were obtained immediately after enrollment on visit day 1 and in follow-up visits on days 6, 14, and 28. Mid turbinate nasopharyngeal swabs (MT-swabs) were collected. Briefly, flocked sterile swabs were inserted approximately 1 inch into the mid-turbinate region, rotated several times against the nasal walls of each nares, and placed into viral transport media (VTM). Nasopharyngeal rinses were next collected. Briefly, both nares were flushed with 0.1% saline and collected by gravity flow or aspiration and transferred to tubes containing ice-cold bronchial epithelial cell growth medium (BEGM) and processed within 2 hours of collection. Rinses were filtered using a 40 µm cell strainer with cells pelleted and removed, then the protease inhibitor cocktail was added to nasopharyngeal rinse fluids (NRF) before they were stored at -80°C. Blood was drawn into BD vacutainer CPT™ cell preparation tubes (BD Biosciences, Franklin Lakes, NJ, USA) with sodium citrate. Blood was processed the day of collection per manufacturer guidelines and plasma stored at -80°C. On study days 14 and 28, participants completed a questionnaire, which included questions to determine estimated dates of symptom onset and relief, socio-demographic and family characteristics, and medical history.

**SARS-CoV-2 RNA RT-qPCR and Sequencing:** MT-swabs were transported to the lab frozen or on ice. RNA was isolated and subjected to reverse transcription real-time quantitative polymerase chain reaction (RT-qPCR) and positive samples sequenced. RNA from MT-swabs were extracted using a RNeasy mini kit (Qiagen, Venlo, Netherlands, #74106) following the manufacturer’s protocol. One-step RT-qPCR reaction was performed with ABI FAST Virus 1-step Master Mix (Applied Biosystems, Waltham, MA, USA, #4444436), SARS-Cov-2 specific primers and probes targeting either ORF1b-nsp14 or Spike (in house design in collaboration with St Jude’s Center for Applied Bioinformatics, CAB) and template RNA on an ABI 7500 FAST machine (Applied Biosystems).(E1) Reverse transcription was performed for 5 minutes at 50°C, followed by 20 seconds at 95°C. The reaction was performed with 40 cycles of 95°C for 5 seconds, followed by 60°C for 30 seconds. A cycle threshold (Ct) value for each sample was determined as the average of triplicate wells per target. Samples failing to cross the threshold by cycle 40 were marked as negative.

SARS-CoV-2 positive MT-swabs samples were sequenced. For sequencing, RNA was isolated from samples using either the Qiagen RNeasy Mini Kit (Qiagen, #74104) or the MagMaxTM Viral/Pathogen Nucleic Acid Isolation Kit (Applied Biosystems, #A42352) on a ThermoFisher Kingfisher Flex 96-well magnetic purification platform (Thermo Fisher Scientific, Waltham, MA, USA). RNA was transcribed to single stranded cDNA using an Invitrogen SuperScript IV first strand synthesis kit (Invitrogen, Waltham, MA, USA, #18091050) using random hexamers and following manufacturer’s recommendations with the minor modification of extending the RT incubation step from 5 minutes to 1 hour. From the cDNA, SARS-CoV-2 sequence libraries were prepared using the xGen SARS-CoV-2 Amplicon Panel (Integrated DNA Technologies [IDT], Coralville, Iowa, USA, #10009832) and following the manufacturer’s protocol. Briefly, the SARS-CoV-2 genome was amplified using a multiplexed PCR creating 345 overlapping amplicons in a single reaction. Adapters and indexes were then added to the amplicons in a second limited cycle PCR. The libraries were normalized using the Normalase enzymatic treatment. Normalized libraries were quantified using the NEBNext Library Quant kit for Illumina (New England Biolabs, Ipswich, MA, USA, #E7630L) and diluted to the required loading concentration. The libraries were sequenced by paired-end 2 x 150 using the Illumina MiSeq Reagent Kit v2 (Illumina, San Diego, CA, USA, #MS-102-2002) for 300-cycles.

The sequenced libraries were assembled, and SARS-CoV-2 lineages were determined using an in-house developed pipeline called idCOV.(E2) Briefly, the reads were trimmed for quality using Trimmomatic and primer sequences were removed using Primerclip (IDT). The reads were then assembled to the original Wuhan-Hu-1 strain sequence. Mutations were identified and confidence was assigned based on read quality and depth at each location. Once the genome was constructed, mutation markers were compared to lineage-defining mutations to determine the SARS-CoV-2 lineage of each sample. Following the idCOV lineage determinations, all lineages were confirmed through UCSC UShER: Ultrafast Sample placement on Existing tRee to compare each assembled genome with all available SARS-CoV-2 sequences.

**Antigen and Antibody Quantification:** SARS-CoV-2 S and N Ag proteins were quantified in plasma and NRF by ELISA (ELV-COVID19S1 and ELV-COVID19N, respectively; RayBiotech, Peachtree Corners, GA, USA). Antibodies (IgM, IgA, and IgG antibodies) against SARS-CoV-2 S and N proteins were quantified in plasma and NRF using ELISA kits from RayBiotech (IEQ-CoVSN-IgM, IEQ-CoVSN-IgA, and IEQ-CoVSN-IgG). Technical replicates were performed. Samples were run in duplicate per manufacturer’s protocol and analyzed per manufacturer recommendations, except that the nasal fluids were not diluted. Antigen and antibody concentrations were extrapolated from standard curves.

**SARS-CoV-2/VSV pseudotype production and neutralization assays:** VSV-DG-luciferase pseudo types displaying the full-length SARS-CoV-2 spike (Wuhan-Hu-1 strain) were generated using a plasmid encoding a codon-optimized cDNA for the SARS-CoV-2 spike as described by Whitt 2010.(E3) Plasmid expressing codon-optimized cDNA for the SARS-CoV-2 spike was generously provided to us by Dr. Florian Krammer, Icahn School of Medicine, Mount Sinai. (E4) Residual infectivity from the VSV-G pseudo typed DG-luciferase inoculum was neutralized immediately after VSV-G pseudo typed DG-luciferase virus adsorption by incubation for 30 minutes with a hybridoma culture supernatant that contained the I1 VSV G-specific monoclonal antibody.(E5) SARS-CoV-2 specific titers were determined on VeroE6-TMPRSS2 cells, which were obtained from the Japanese Collection of Research Bioresources (JCRB) (National Institutes of Biomedical Innovation, Health and Nutrition, Japan, Cat #1819).(E6)

SARS-CoV-2 neutralizing activity in sera from study participants was determined in VeroE6-TMPRSS2 cells using a standard protocol. Briefly, 2 x 103 infectious units of SARS-CoV-2/DG-luciferase pseudotype virus were mixed with increasing 2-fold dilutions of patient sera that had been complement-inactivated by incubation at 56°C for 30 minutes. The virus-sera admixtures were next incubated at 37°C for 1 hour and then added directly to VeroE6-TMPRSS2 cells in a 96-well plate that had been passaged ~18 hours prior. Luciferase activity was assayed 17 hours post-inoculation using the XTND Luc-Screen kit as per the manufacturer’s instructions (Applied Biosystems) and relative light units were read using a BioTek Synergy 2 plate reader (BioTek, Winooski, Vermont, USA). The raw data were then transformed and plotted using Prism GraphPad (GraphPad Software, La Jolla, CA, USA) and neutralizing titers were determined and compared to a positive control (sera from an individual who had severe COVID-19) and a negative control serum from an uninfected individual. Ab neutralization potency was calculated as a ratio of neutralization titer 50 (NT_50_) to the sum of plasma SARS-CoV-2 specific isotypes and grouped by severity.(E7)

**Bio-Plex Neutralization Antibody Assay:** Plasma from study participants collected on either study day 28 or 14 days after the diagnosis was quantified for the percent spike and RBD antibody inhibition using Bio-Plex Pro Human SARS-CoV-2 neutralization antibody assay kit from Bio-Rad (Bio-Rad Laboratories, Hercules, CA, USA, #12016848). Neutralization antibody assay with each sample was run in duplicate and the assay was performed according to the manufacturer’s instructions and then plate was analyzed on a Luminex 200 (Luminex Multiplexing Instrument, Merck Millipore, Burlington, MA, USA) analyzer. Data obtained were extracted using xPONENT software (IQVIA, Durham, NC, USA). Percent inhibition was calculated by comparing sample mean fluorescent intensity (MFI) results to the negative control using the following formula: percentage inhibition = (1 – sample MFI / MFI negative control) x 100.

**Mucosal and Plasma Cytokine and Chemokine Quantification:** Longitudinal frozen plasma and nasal fluids were analyzed for 41 cytokines/chemokines levels using a human cytokine/chemokine magnetic bead panel HCYTMAG-60K-PX41 Millipore Sigma (Merck Millipore). Nasal fluid was concentrated before the assay and total protein quantified for normalization across biospecimens. Cytokines, chemokines, and growth factors were quantified using on a Luminex 200 Multiplexing Instrument (Luminex, Merck Millipore). The Luminex assay was performed according to the manufacturer’s instructions using duplicate technical replicates. Data obtained were extracted using xPONENT software. The Absolute quantity of cytokines detected was reported as pg/mL of plasma or as pg/mg protein of nasal fluid.

**Statistical Analysis:** The demographic, disease symptoms, comorbidities and treatment characteristics of study participants were described using the number and percent of available data for participants in the study by COVID-19 severity. The age of the cohort described using the median and interquartile range (defined by the 75^th^ percentile). GraphPad Prism software was used for basic statistical analysis: two-tailed Mann–Whitney U test to determine mean differences in viral load between weeks 1 and ≥2; Log-rank Mantel-Cox test for Ab conversion rates, one-way ANOVA with Fisher’s least significant difference (LSD) procedure, followed by a posttest for linear trends for mean differences in Ab magnitude, and two-way ANOVA with Tukey's honestly significant difference (HSD) test for mean differences in cytokine expression among outcome groups over time. XLSTAT package was used for expression analysis, heatmaps, correlation analysis, principal component analysis (PCA), partial least squares-discriminant analysis (PLS-DA), and Classification and Regression Tree (CART) analysis. Correlations between the different parameters were calculated using the Pearson’s correlation test. Asterisks symbolize p values from these analyses as follows: *P* < 0·05 (*), *P* < 0·01 (**), *P* < 0·001 (***), and *P* < 0·0001 (****). Correlations between the different parameters were calculated using the Pearson’s correlation test in XLSTAT and GraphPad software. The significantly correlated parameters in both the software (XLSTAT and GraphPad) were plotted using GraphPad. *P* values from these analyses are presented as a single asterisk (*) for *P* < .05, two asterisks (**) for *P* < .01, three asterisks (***) for *P* < .001, and four asterisks (****) for *P* < .0001.

**Table E1. Antibody phenotypes and concentrations by hierarchical clusters.**

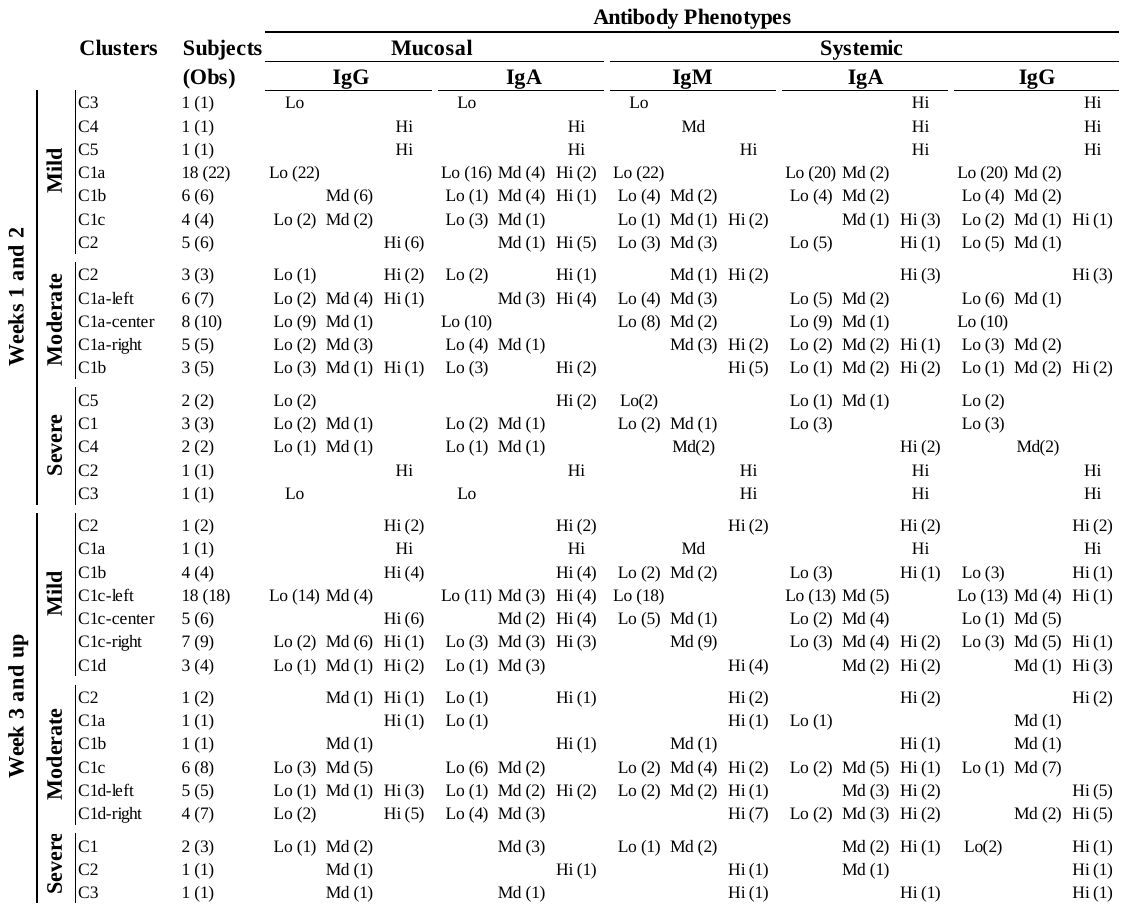

Subjects and paired observations (Obs) per cluster; Low (Lo) < Quartile 2; Middle range (Md) > Quartile 2 and < Quartile 3; High (Hi) ≥ Quartile 3.

**Table E2. Significant difference between children and adults with moderate and severe COVID-19**

| **Variables** | ***P*-value** | **Adult pg/ml** | **Child pg/ml** | **Class** | **Type** | **Response** | **Main Source** |
| --- | --- | --- | --- | --- | --- | --- | --- |
| S-Ag (p) | **<0.0001** | 7,722 (b) | 82,181 (a) | N/A | Antigen | N/A | Virus |
| sCD40L (p) | **0.001** | 182,391 (b) | 377,244 (a) | Adaptive | Ligand | Pro-inflammatory | PLT^a^, T-cells^a^, MØ |
| Fractalkine (p) | **0.001** | 78,409 (b) | 141,486 (a) | Both | GF | Pro-inflammatory | EC |
| G-CSF (p) | **0.001** | 50,69 (b) | 79,446 (a) | Innate | GF | Pro-inflammatory | EC, MØ, Mono |
| IL-1RA (p) | **0.008** | 48,64 (b) | 68,42 (a) | Innate | Cytokine | Anti-inflammatory | Neut, MØ, Mono |
| MCP-3 (p) | **0.040** | 60,067 (b) | 90,473 (a) | Both | Chemokine | Pro-inflammatory | MØ |
| IFN-α2 (n) | **0.046** | 24,803 (a) | 1,766 (b) | Innate | Cytokine | Pro-inflammatory | DC, Neut, MØ |
| S-nAb (p) | **0.049** | 67,245 (a)* | 52,695 (b)* | Adaptive | Antibody | Humoral | B-cells |
| Fractalkine (n) | **0.049** | 2,343,549 (a) | 642,096 (b) | Both | GF | Pro-inflammatory | EC |
| IL-4 (p) | 0.065 | 275,957 (a) | 499,711 (a) | Adaptive | Cytokine | Pro-inflammatory | T-cells, Neut |
| IgA (p) | 0.074 | 18,490,752 (a) | 35,780,315 (a) | Adaptive | Antibody | Humoral | B-cells |
| IL-1β (n) | 0.080 | 14,202 (a) | 52,892 (a) | Innate | Cytokine | Pro-inflammatory | MØ |
| MDC (n) | 0.082 | 40,518 (a) | 60,545 (a) | Innate | Chemokine | Pro-inflammatory | MØ, DC |

Adult n=22 (study days n=72), Child n=17 (study days n=32); Plasma (p), Nasal Rinse Fluids (n), Spike neutralizing Ab titer (S-nAb), Growth factor (GF), Activated (^a^), Platelets (PLT), Macrophages (MØ), Endo/Epithelial cells (EC), Neutrophils (Neut), Monocytes (Mono), Dendritic Cells (DC); *nAb titers are relative not absolute pg/ml.

**Table E3. Classification and Regression Tree structure and rules for COVID-19 severity predictions irrespective of time from diagnosis**

| **Node** | **VO** | **PO (%)** | **Split**  **Variable** | **Rule** | **Predicted Value** | **Node**  **Purity (%)** |
| --- | --- | --- | --- | --- | --- | --- |
| 1 | 233 | 100 |  |  | Mild | 44 |
| 2 | 80 | 34 | IL-1RA (P) | If IL-1RA (P) ≤ 19.25 then Severity = Mild in 69% of cases | Mild | 89 |
| 3 | 153 | 66 | IL-1RA(P) | If IL-1RA (P) > 19.25 then Severity = Moderate in 94% of cases | Moderate | 49 |
| 4 | 73 | 31 | MDC (P) | If IL-1RA (P) ≤ 19.25 and MDC (P) ≤ 595.48 then Severity = Mild in 64% of cases | Mild | 90 |
| 5 | 7 | 3 | MDC (P) | If IL-1RA (P) ≤ 19.25 and MDC (P) > 595.48 then Severity = Mild in 5% of cases | Mild | 71 |
| 6 | 134 | 58 | IL-8 (N) | If IL-1RA (P) > 19.25 and IL-8 (N) ≤ 18,736.73 then Severity = Moderate in 94% of cases | Moderate | 56 |
| 7 | 19 | 8 | IL-8 (N) | If IL-1RA (P) > 19.25 and IL-8 (N) > 18,736.73 then Severity = Severe 38% of cases | Severe | 100 |
| 8 | 69 | 30 | TGFα (P) | If IL-1RA (P) ≤ 19.25 and MDC (P) ≤ 595.48 and TGFα (P) ≤ 2.10 then Severity = Mild in 63% of cases | Mild | 94 |
| 9 | 4 | 2 | TGFα (P) | If IL-1RA (P) ≤ 19.25 and MDC (P) ≤ 595.48 and TGFα (P) > 2.10 then Severity = Severe in 4% of cases | Severe | 50 |
| 12 | 43 | 19 | TNFβ (P) | If IL-1RA (P) > 19.25 and IL-8 (N) ≤ 18,736.73 and TNFβ (P) ≤ 8.07 then Severity = Severe in 40% of cases | Severe | 47 |
| 13 | 91 | 39 | TNFβ (P) | If IL-1RA (P) > 19.25 and IL-8 (N) ≤ 18,736.73 and TNFβ (P) > 8.07 then Severity = Moderate in 71% of cases | Moderate | 63 |
| 16 | 49 | 21 | IgM (P) | If IL-1RA (P) ≤ 19.25 and MDC (P) ≤ 595.48 and TGFα (P) ≤ 2.10 and IgM (P) ≤ 20,435.21 then Severity = Mild in 48% of cases | Mild | 100 |
| 17 | 20 | 8.6 | IgM (P) | If IL-1RA (P) ≤ 19.25 and MDC (P) ≤ 595.48 and TGFα (P) ≤ 2.10 and IgM (P) > 20,435.21 then Severity = Mild in 16% of cases | Mild | 80 |
| 24 | 21 | 9 | IL-3 (N) | If IL-1RA (P) > 19.25 and IL-8 (N) ≤ 18,736.73 and TNFβ (P) ≤ 8.07 and IL-3 (N) ≤ 8.51 then Severity = Moderate in 18% of cases | Moderate | 67 |
| 25 | 22 | 9 | IL-3 (N) | If IL-1RA (P) > 19.25 and IL-8 (N) ≤ 18,736.73 and TNFβ (P) ≤ 8.07 and IL-3 (N) > 8.51 then Severity = Severe in 32% of cases | Severe | 73 |
| 26 | 71 | 30 | IFNγ (P) | If IL-1RA (P) > 19.25 and IL-8 (N) ≤ 18,736.73 and TNFβ (P) > 8.07 and IFNγ (P) ≤ 25.50 then Severity = Moderate in 61% of cases | Moderate | 69 |
| 27 | 20 | 9 | IFNγ (P) | If IL-1RA (P) > 19.25 and IL-8 (N) ≤ 18,736.73 and TNFβ (P) > 8.07 and IFNγ (P) > 25.50 then Severity = Mild in 12% of cases | Mild | 60 |
| Number of observations the rule is verified by (VO); Percent of observations in the node (PO); Plasma (P); Nasal Fluid (N); Percent of the node that belong to the dominate dependent variable (Node Purity) | | | | | | |

**Table E4. Classification and Regression Tree structure and rules for COVID-19 severity predictions in acute phase (within 1 week of diagnosis)**

| **Node** | **VO** | **PO (%)** | **Split**  **Variable** | **Rule** | **Predicted Value** | **Node**  **Purity (%)** |
| --- | --- | --- | --- | --- | --- | --- |
| 1 | 80 | 100 |  |  | Moderate | 39 |
| 2 | 68 | 85 |  | If IL-1β (N) ≤ 29.74 then Severity= Moderate in 100% of cases | Moderate | 46 |
| 3 | 12 | 15 | IL-1β (N) | If IL-1β (N) > 29.74 then Severity= Severe in 52% of cases | Severe | 100 |
| 4 | 15 | 19 | IL-1β (N) | If IL-1β (N) ≤ 29.74 and IL-1RA (P) ≤ 15.46 then Severity= Mild in 54% of cases | Mild | 93 |
| 5 | 53 | 66 | IL-1RA (P) | If IL-1β (N) ≤ 29.74 and IL-1RA (P) > 15.46 then Severity= Moderate in 97% of cases | Moderate | 57 |
| 8 | 13 | 16 | IL-1RA (P) | If IL-1β (N) ≤ 29.74 and IL-1RA (P) ≤ 15.46 and IFNy (N) ≤ 10.33 then Severity= Mild in 50% of cases | Mild | 100 |
| 9 | 2 | 3 | IFNy (N) | If IL-1β (N) ≤ 29.74 and IL-1RA (P) ≤ 15.46 and IFNy (N) > 10.33 then Severity= Mild in 4% of cases | Mild | 50 |
| 10 | 48 | 60 | IFNy (N) | If IL-1β (N) ≤ 29.74 and IL-1RA (P) > 15.46 and IgA (P) ≤ 156,785.92 then Severity= Moderate in 97% of cases | Moderate | 63 |
| 11 | 5 | 6 | IgA (P) | If IL-1β (N) ≤ 29.74 and IL-1RA (P) > 15.46 and IgA (P) > 156,785.92 then Severity= Severe in 22% of cases | Severe | 100 |
| 20 | 35 | 44 | IgA (P) | If IL-1β (N) ≤ 29.74 and IL-1RA (P) > 15.46 and IgA (P) ≤ 156,785.92 and CD40L (N) ≤ 27.78 then Severity= Moderate in 81% of cases | Moderate | 71 |
| 21 | 13 | 16 | CD40L (N) | If IL-1β (N) ≤ 29.74 and IL-1RA (P) > 15.46 and IgA (P) ≤ 156,785.92 and CD40L (N) > 27.78 then Severity= Severe in 26% of cases | Severe | 46 |
| Number of observations the rule is verified by (VO); Percent of observations (PO); Plasma (P); Nasal Fluid (N); Percent of the node that belong to the dominate dependent variable (Node Purity) | | | | | | |

**Supplementary Figures Legends:**

**Figure E1. Study design and longitudinal samples collected. A** Schematic representation of biospecimens (Blood, nasal rinses, and nasal swabs) collected over one month (study days 1, 6, 14, and 28) from 78 COVID-19 positive subjects (mild [n = 27], moderate [n = 29], and severe [n = 22]) with 42% males and 58% females. **B** The total number of biospecimens collected at different study day grouped by disease severity. Total number of biospecimens collected longitudinally includes MT-swabs from 78, 28, 27, and 45 participants, blood from 69, 44, 40, and 39 participants and NRF from 75, 44, 46, and 44 participants on study days 1, 6, 14, and 28 respectively.

**Figure E2. SARS-CoV-2 viral and infection stage characterization.** Viral RNA measured in swabs using quantitative real time RT-PCR (qRT-PCR) in terms of Ct value and viral antigen in NRF and plasma by ELISA. Each circle represents a patient colored by severity; mild (
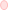
), moderate (●), and severe (●). Positivity thresholds are represented with dotted lines. **A** Viral RNA in nasal swabs over time (left) and between paired swabs at acute and recovery phase (right). Viral RNA was isolated from swabs and quantified by qRT-PCR using SARS-CoV-2 specific primers. Ct values were plotted by time from diagnosis. Cycle threshold (Ct) was 40 and PCR positive samples plotted by severity groups. Ct values for subjects with mild, moderate, and severe COVID-19 were grouped into the acute and recovery phases (i.e., days 1–7 and >7 days from diagnosis, respectively) and were compared by paired student’s t-test (****P* ≤ .001, **** *P* ≤ .0001, and ns: non-significant). Two of severe patients died during the study and indicated as star (**_*_**) in the plot. **B** SARS-CoV-2 specific nucleocapsid and spike proteins in nasal rinses (left) and in plasma (right) respectively and grouped by disease severity. Severity groups were compared by one-way ANOVA with Tukey's HSD test (*P < .05 and ** P < .01). **C** Viral RNA sequencing data from PCR positive swabs (n = 52) plotted by month and percent of SARS-CoV-2 alpha sub-variants each month from July 2020 – Jan 2021. **D** Severe COVID-19 patients -dominate Ag-specific IgM+IgG+ sero-phenotype within a week of SARS-CoV-2 infection. Percent of IgM-IgG-, IgM+IgG- and IgM+IgG+ SARS-CoV-2 specific antibody phenotype in patients were grouped by disease severity and duration (weeks).

**Figure E3. S and N-specific antibody kinetics in mucosa and periphery.** Ag-specific antibodies in NRF and plasma measured by ELISA over time. Each circle represents a patient colored by COVID-19 severity groups (mild [
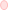
], moderate [●], and severe [●]). **A, B** Ag-specific Ab kinetics in NRF (**A**) and in plasma (**B**). Dotted line represents the cut-off of positivity (neg), mid Ab levels (Q2) and high Ab levels (Q3). Between neg and Q2 are participants showing low Ab levels, Q2 and Q3 are participants showing mid Ab levels and above Q3 are participants showing high Ab levels.

**Figure E4. Mucosal and systemic S and N-specific antibody trends by outcome severity and by week from diagnosis.** Antibodies were quantitated in the NRF collected longitudinally from participants with mild (n = 102 samples), moderate (n = 75 samples), and severe (n = 27 samples) disease severity and in the plasma samples from participants with mild (n = 86 samples), moderate (n = 59 samples), and severe (n = 26 samples) disease severity using an ELISA. Dotted line represents cut-off of positivity. Each circle represents a patient colored by COVID-19 severity groups (mild [
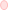
], moderate [●], and severe [●]). **A, B** Linear trend analysis of mucosal IgA and IgG among severity groups (**A**) and within a group (**B**) overtime. **C, D** Linear trend analysis of plasma IgA, IgG and IgM among severity groups (**C**) and within a group (**D**) overtime. Longitudinal samples were grouped by weeks from COVID-19 clinical diagnostic test. Slopes with significant P value are inset in the respective graphs.

**Figure E5. Mucosal and systemic Antigen-specific antibody correlations. A** Correlation plots for paired variables correlated in mild COVID-19 illness. Patients with significant correlation *P* value in XLSTAT software were re-analyzed in GraphPad and correlation coefficient with significant *P* value plotted individually. Pearson's correlation coefficient with significant *P* values (**P* <. 05, *** *P* ≤ .001 and **** *P* ≤ .0001) indicated on the respective plot. Non-paired observations (either NRF or plasma samples missing for the respective study day) were removed from the analysis. Significant antibody correlation was not observed by GraphPad analysis for moderate and severe COVID-19 patients. **B, C** Mucosal and systemic Ag-specific antibody levels in paired longitudinal samples from 0–14 days (**B**) and after 14 days from days from diagnosis (**C**) by severity. Paired NRF and plasma samples with 80 and 72 observations within 2 weeks and 3 or more weeks from diagnosis, respectively. Within each severity group, subjects were grouped by antibody levels using agglomerative hierarchical clustering using Euclidean distance and Ward’s method. Dissimilarity scores of clusters are indicated with major clusters colorized and subcluster labeled alphabetically from left to right on dendrograms. Minimum and maximum antibody levels are reflected in the permuted matrix with Ab pg/ml data values replaced by corresponding color intensities from dark blue to yellow (-1 to 1, respectively).

**Figure E6. Mucosal cytokine quantification and analysis.** Immune factors were quantified in the NRF samples collected longitudinally (n = 55, 33, 33, and 29 subjects from study day 1, 6, 14, and 28, respectively) on the Luminex® xMAP™ system with a MILLIPLEX MAP human cytokine/chemokine immunoassay. Study day was converted to time from positive diagnosis. **A** Immune factors were grouped by major immune functions: anti-inflammatory (orange), adaptive immunity (blue), pro inflammatory (purple), growth factors (red), and chemoattractants (green). **B** Heat maps of statistically different (P < .05) cytokines between COVID-19 participants with mild severity and moderate to severe disease severity, ordered by hierarchical clustering using XLSTAT software. Upregulated cytokines are shown in yellow and downregulated in blue (-1 to 1). **C** Volcano plot of differentially expressed cytokines (-log10 P value > 2 and log2 fold change ≥ 2) in mucosa of mild COVID-19 cases compared to severe. Cytokines that were significantly increased or decreased more than two-log-fold are shown in yellow and blue color, respectively. Each colored filled circle/dot represents an immune factor on the volcano plot. **D–G** Cytokines identified in expression analysis and grouped as chemoattractant (**D**), Growth factors (**E**), proinflammatory cytokines (**F**) and anti-inflammatory cytokines (**G**). Severity groups, mild (
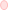
), moderate (●), and severe (●) and weeks were compared using two-way ANOVA with Tukey's HSD test (**P* < .05, ** *P* < .01, *** *P* < .001 and **** *P* < .0001).

**Figure E7. Systemic cytokine quantification and analysis**. Immune factors were quantified in plasma samples collected longitudinally from mild (n = 90 samples), moderate (n = 56 samples), and severe (n = 27 samples) cases on the Luminex® xMAP™ system with a MILLIPLEX MAP human cytokine/chemokine immunoassay. **A** Immune factors were grouped by major immune functions: anti-inflammatory (orange), adaptive immunity (blue), pro inflammatory (purple), growth factors (red), and chemoattractants (green). **B** Heat maps of statistically different (P < .05) cytokines between COVID-19 participants with mild severity and moderate to severe disease severity, ordered by hierarchical clustering using XLSTAT software. Upregulated cytokines are shown in yellow and downregulated in blue (-1 to 1). **C** Volcano plot of differentially expressed cytokines (-log10 *P* value > 2 and log2 fold change ≥ 2) in plasma of mild COVID-19 participants compared to moderate and severe. Plasma cytokines that were significantly increased or decreased more than two-log-fold in moderate to severe participants compared to mild are shown in yellow and blue color, respectively. Each colored filled circle/dot represents an immune factor on the volcano plot. **D** Individual cytokine magnitudes expression plots by COVID-19 severity longitudinally (IL-1RA, IL-8, IL-6, IL-10, IP-10, IL-15, MIP-1β, Fractalkine, and GM-CSF). Plasma collected from each participant at least at one study day and maximum for four study day (1, 6, 14, and 28). The time from diagnosis was calculated by adding the interval time from diagnostic to the study day. Severity groups (mild [
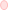
], moderate [●], and severe [●]) and weeks were compared by two-way ANOVA with Tukey's HSD test (**P* < .05, ** *P* ≤ .01, *** *P* ≤ .001 and **** *P* ≤.

**Figure E8. Associations between qualitative and quantitative variables.** Partial least square discriminant analysis (PLS-DA) was performed to further distinguish COVID-19 severity groups and identify qualitative and quantitative parameters contributing to the group separation. **A** Correlations on t1 and t2. Only immune factor variables > |0.4| are labeled. **B** Observations on t1 and t2 clustered into mild (
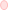
), moderate (●), and severe (●), and moderate cluster overlapping with mild and severe observations. **C** Variable Importance for the Projection (VIP) plotted for variables with VIP score above 1 in at least one of the three models. Variables detected in NRF and plasma are symbolized with n and p respectively, right to the biomarker name in the brackets.

**Figure E9. Immune predictors for mild, moderate, or severe COVID-19.** Supervised machine learning was performed with XLSTAT using the Classification and Regression Tree algorithm including all studied parameters. Parameters were set to minimum parent node size of 5, minimum son node size of 2, maximum tree depth of 3(A) or 4 (B), and complexity parameter of 0.001. Pie diagrams at each node is colored with percent purity of node for severity groups mild (
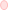
), moderate (●), and severe (●). **A, B** Immune predictors of COVID-19 severity, irrespective of time from diagnosis (**A**) and during acute-phase (**B**). The model included subjects with mild (n=27), moderate (n=29), and severe (n=22) study participants and experimental data from all study visits for mild (n=103), moderate (n=80), and severe (n=50). The total percentage of well-classified observations was 78% (80% for mild, 79% for moderate, and 74% for severe). For acute phase analysis model was restricted to week 1 from diagnosis for mild (n=26), moderate (n=31), and severe (n=23). The total percentage of well-classified observations was 78% (54% for mild, 81% for moderate, and 100% for severe). PCR Ct values and mucosal and systemic immune factors (quantitative variables) and week from diagnosis and antigen positivity (qualitative variables) data were input for subjects with mild (n=27), moderate (n=29), and severe (n=22) COVID-19 (dependent variables). The decision tree branch splits at a predictive variable and is capped by a node. Antibody and cytokine cutoff levels above split variable in AU/ml and pg/ml, respectively, with the node number, size (observations per node), and purity (%). COVID-19 severity groups The purity indicates the percentage of objects that belong to the dominant severity category at each node. Biomarkers in NRF and plasma are symbolized with N and P respectively, right to the biomarker name in the brackets.

**Supplemental Figures**

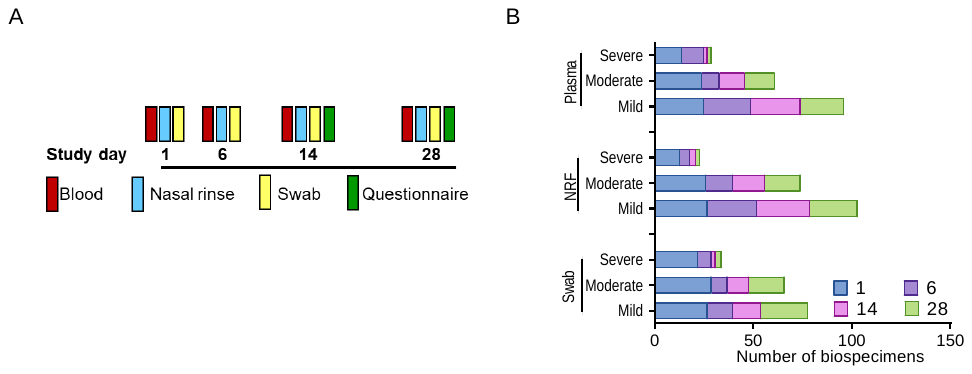

**Figure E1.** Study design and longitudinal samples collected.

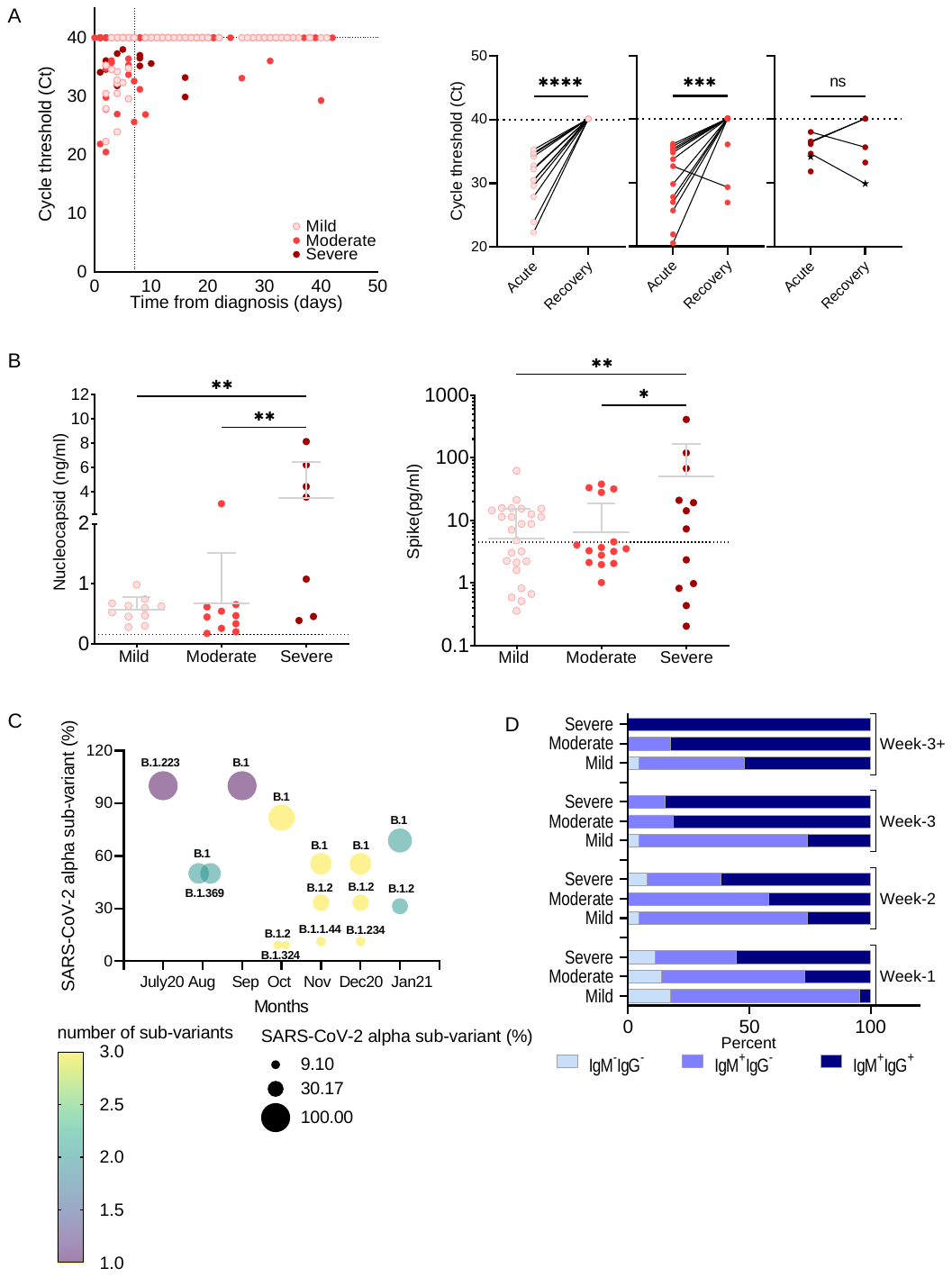
**Figure E2.** SARS-CoV-2 viral and infection stage characterization.

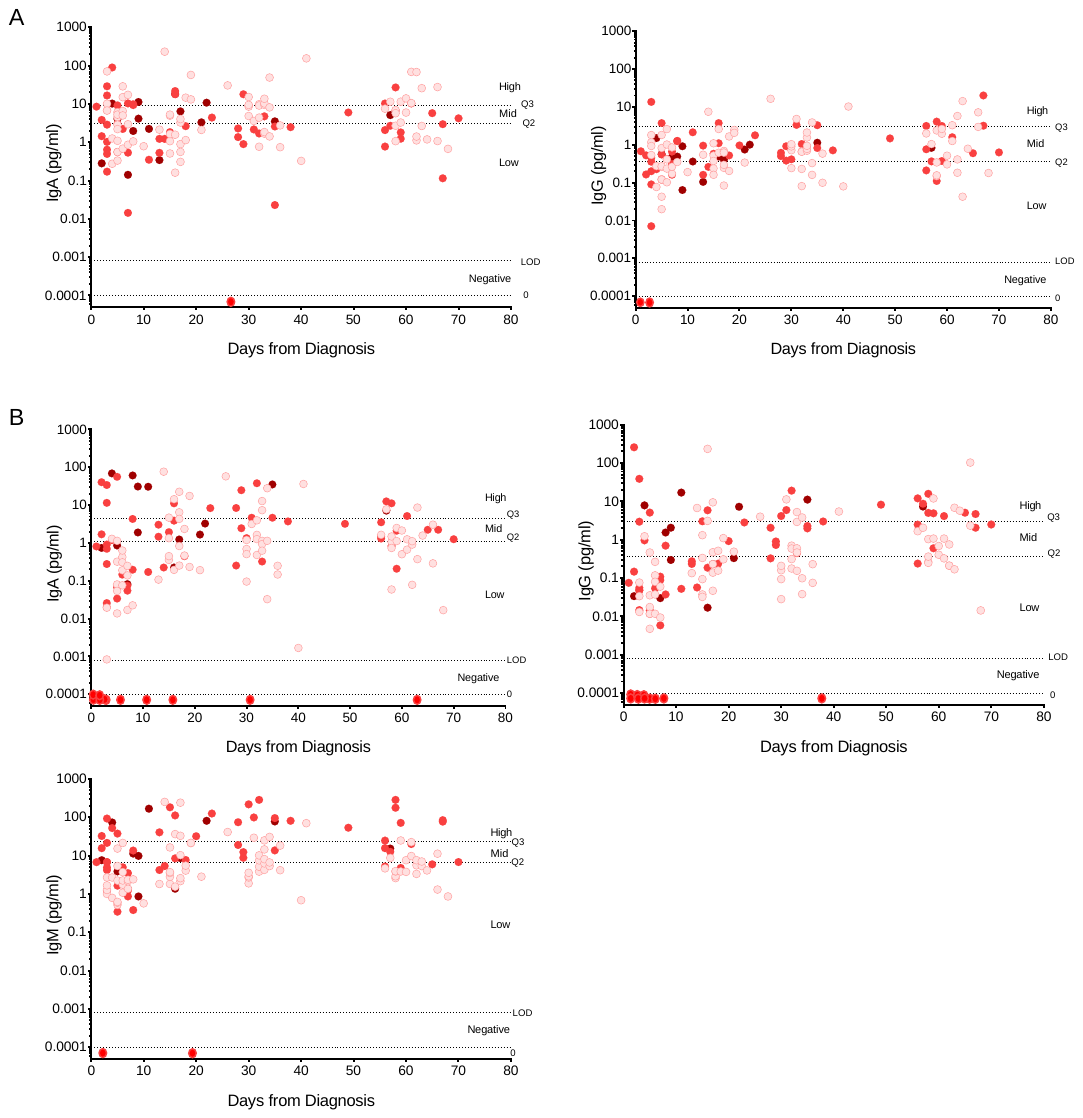
**Figure E3.** S and N-specific antibody kinetics in mucosa and periphery.

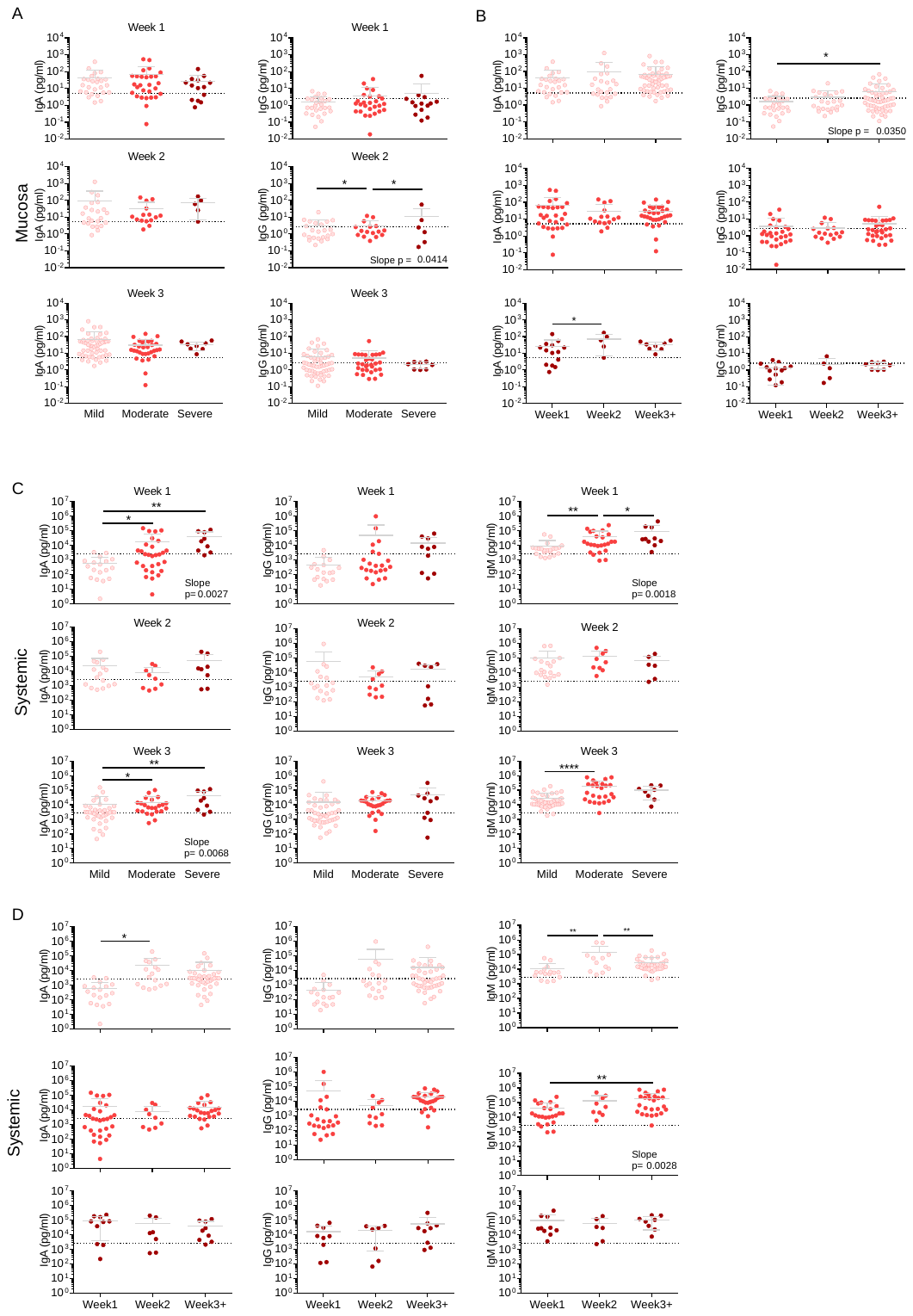
 **Figure E4.** Mucosal and systemic S and N-specific antibody trends by outcome severity and by week from diagnosis.

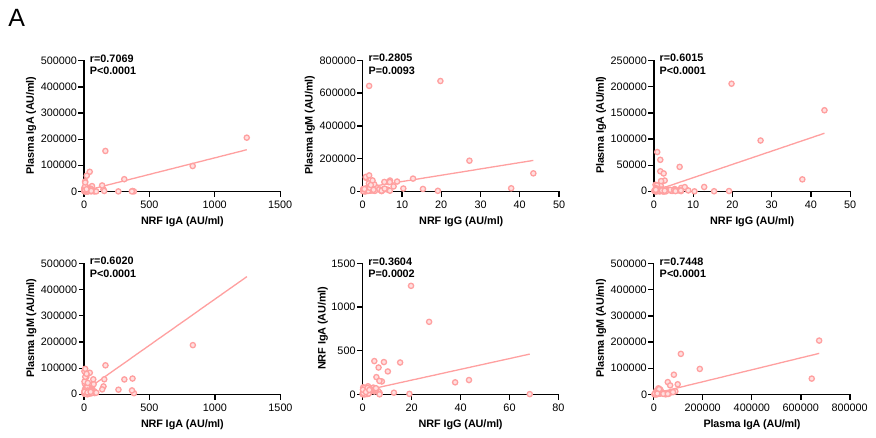

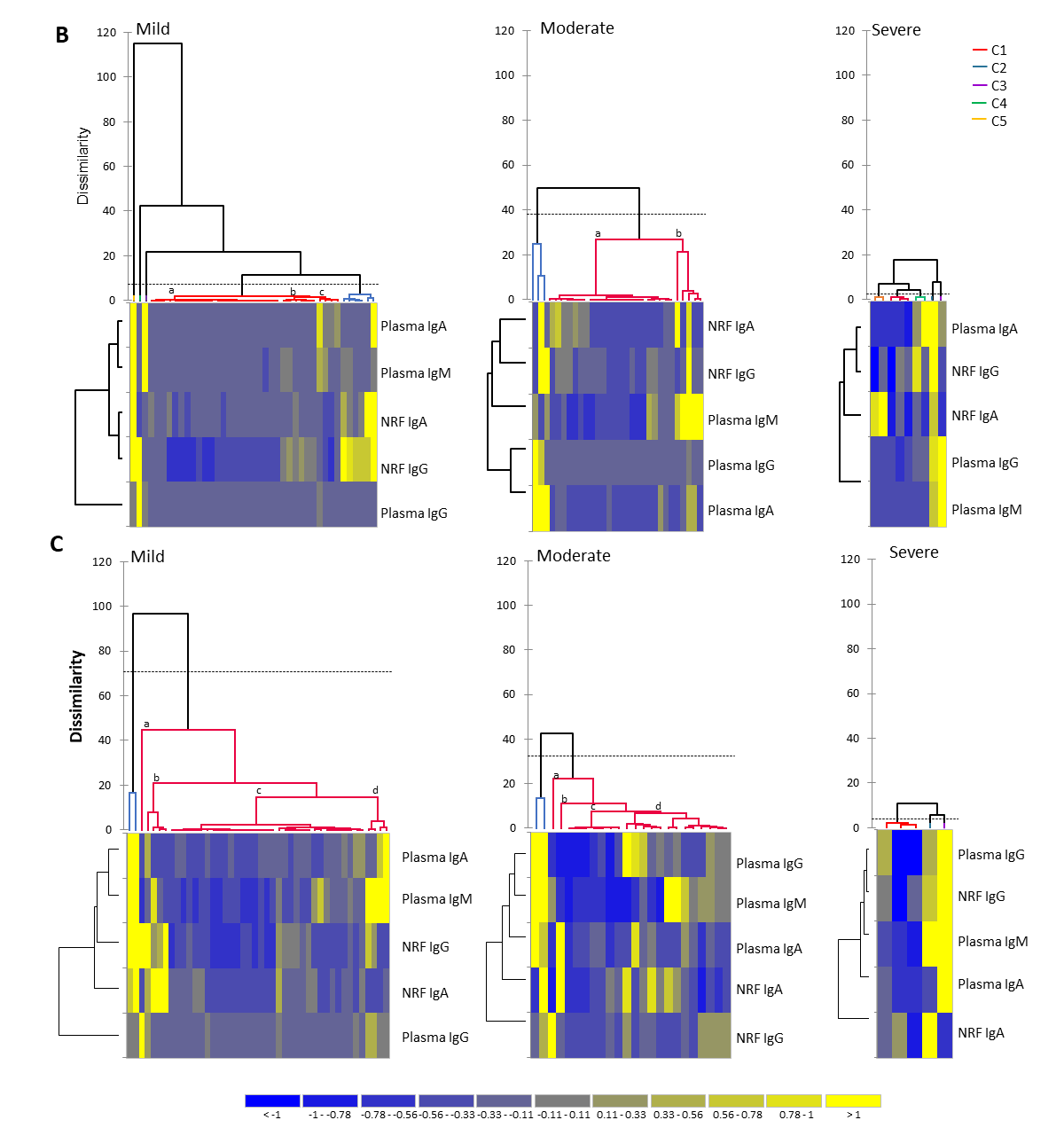

**Figure E5.** Mucosal and systemic Antigen-specific antibody correlations.

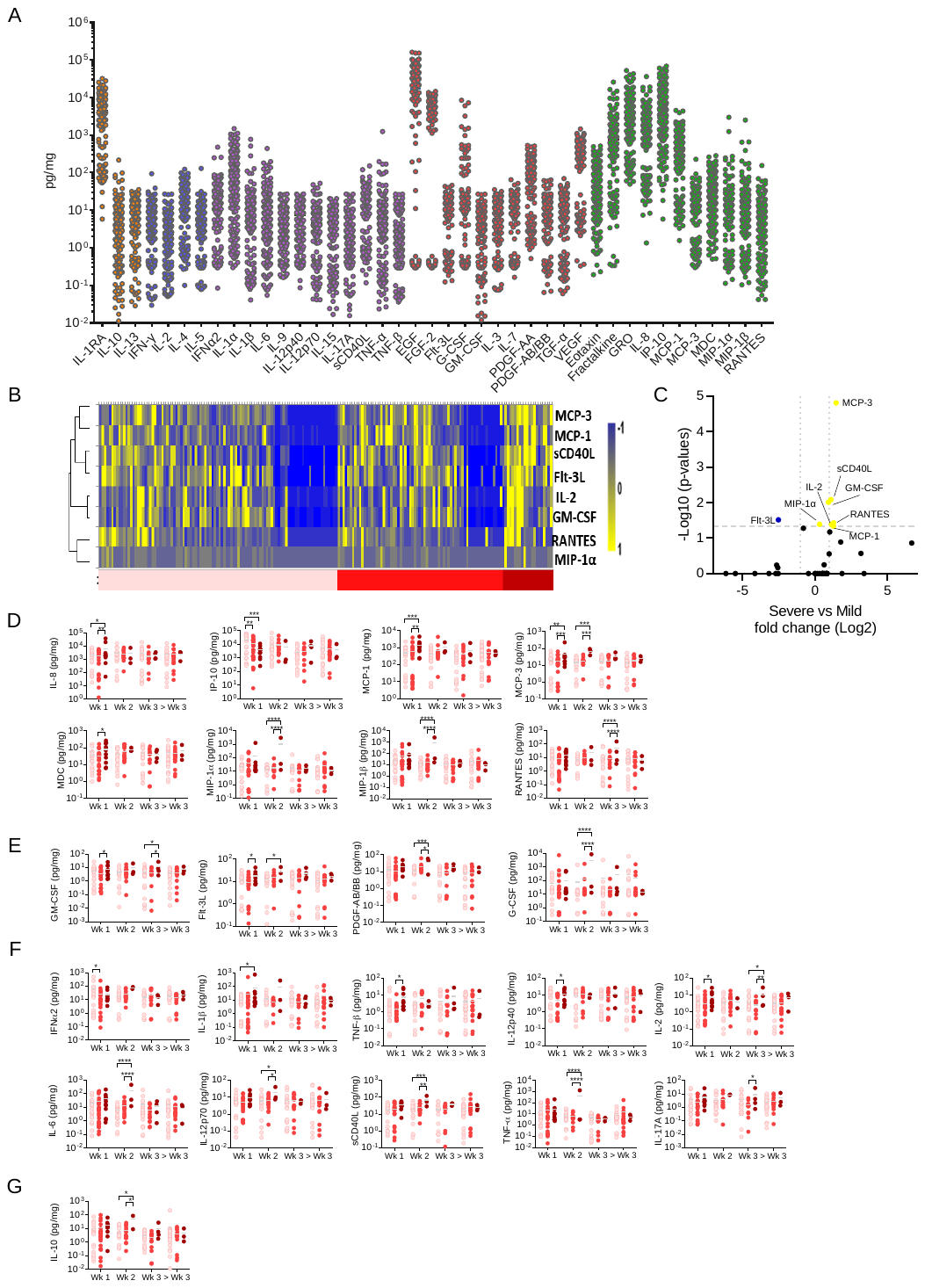
 **Figure E6.** Mucosal cytokine quantification and analysis.

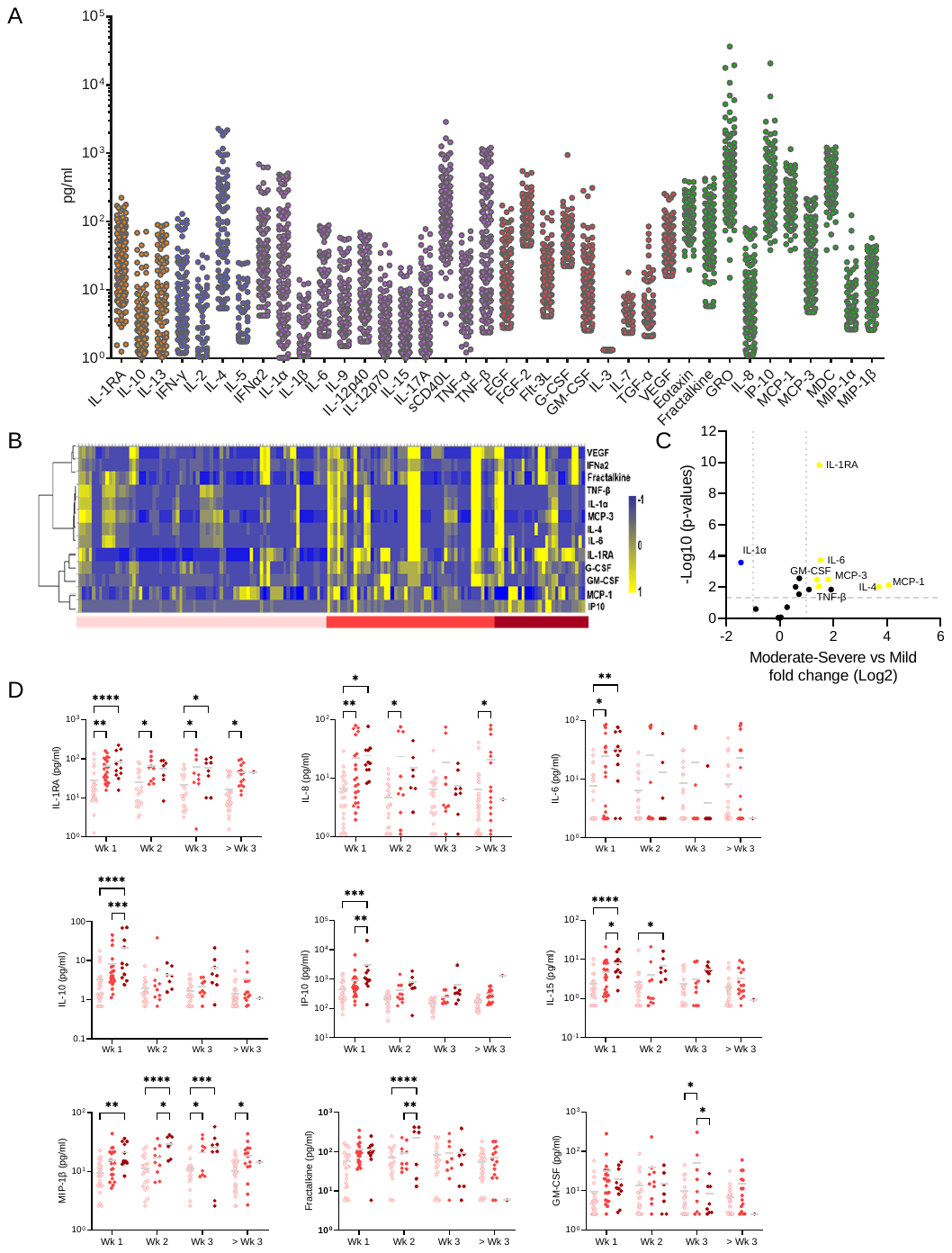

**Figure E7.** Systemic cytokine quantification and analysis.

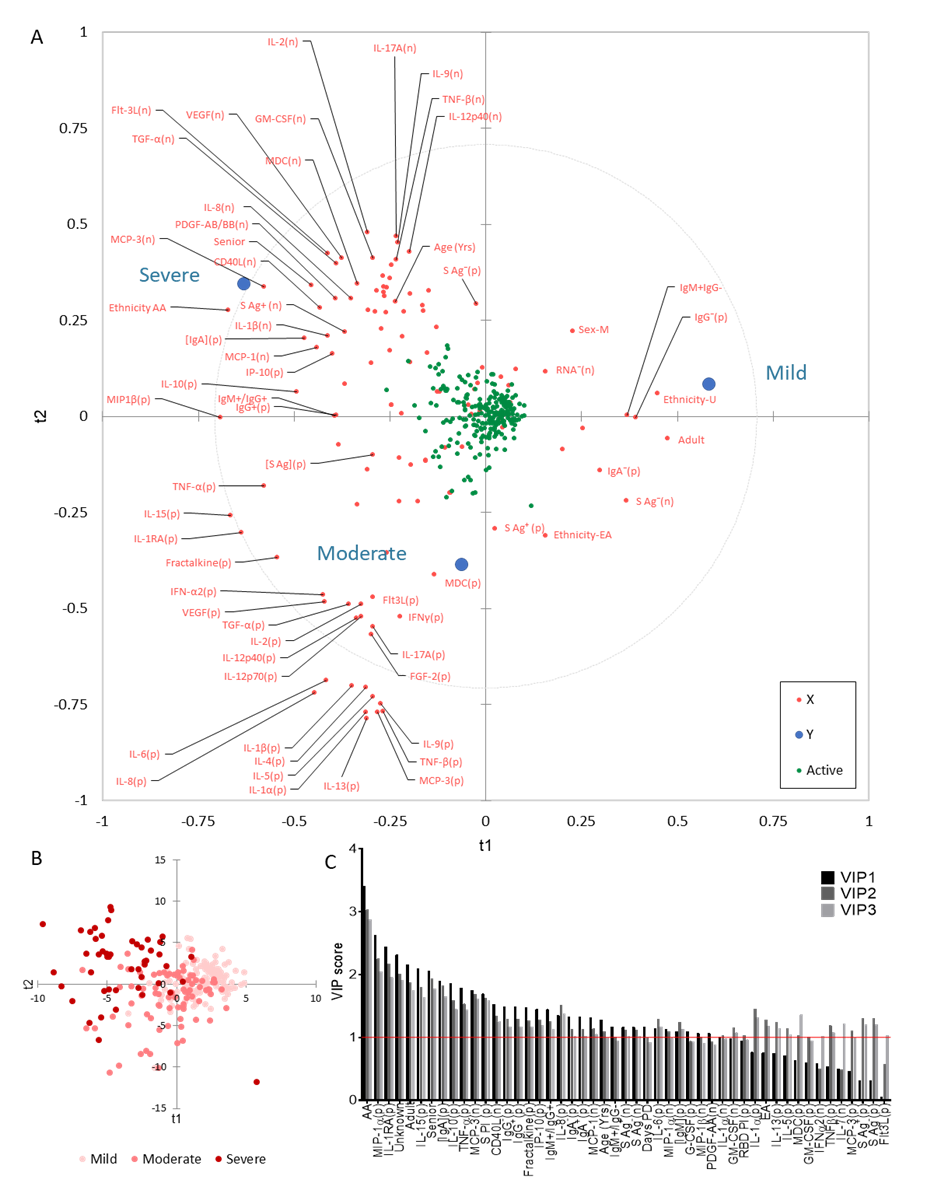

**Figure E8.** Associations between qualitative and quantitative variables.

A

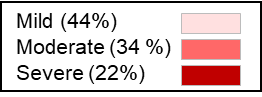

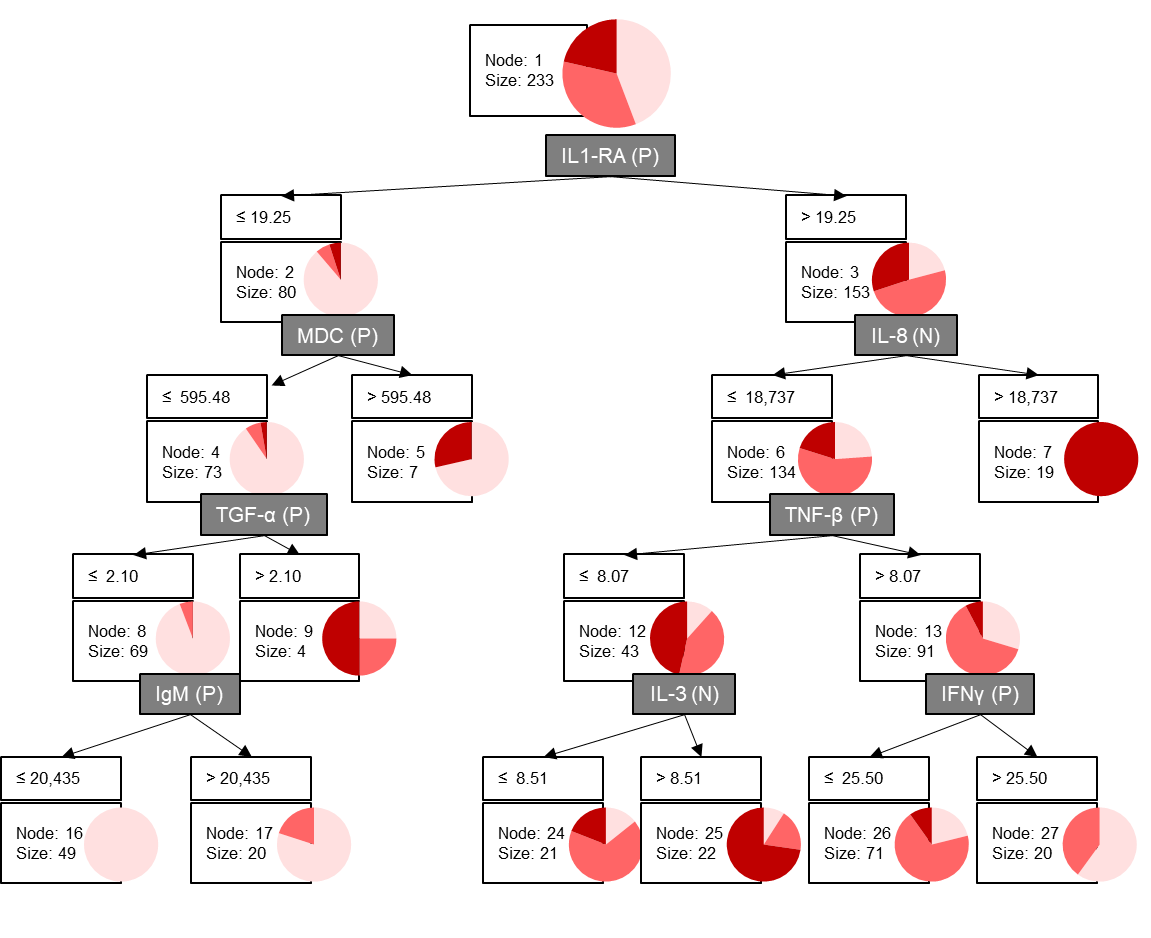

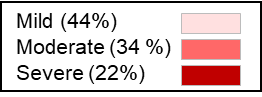

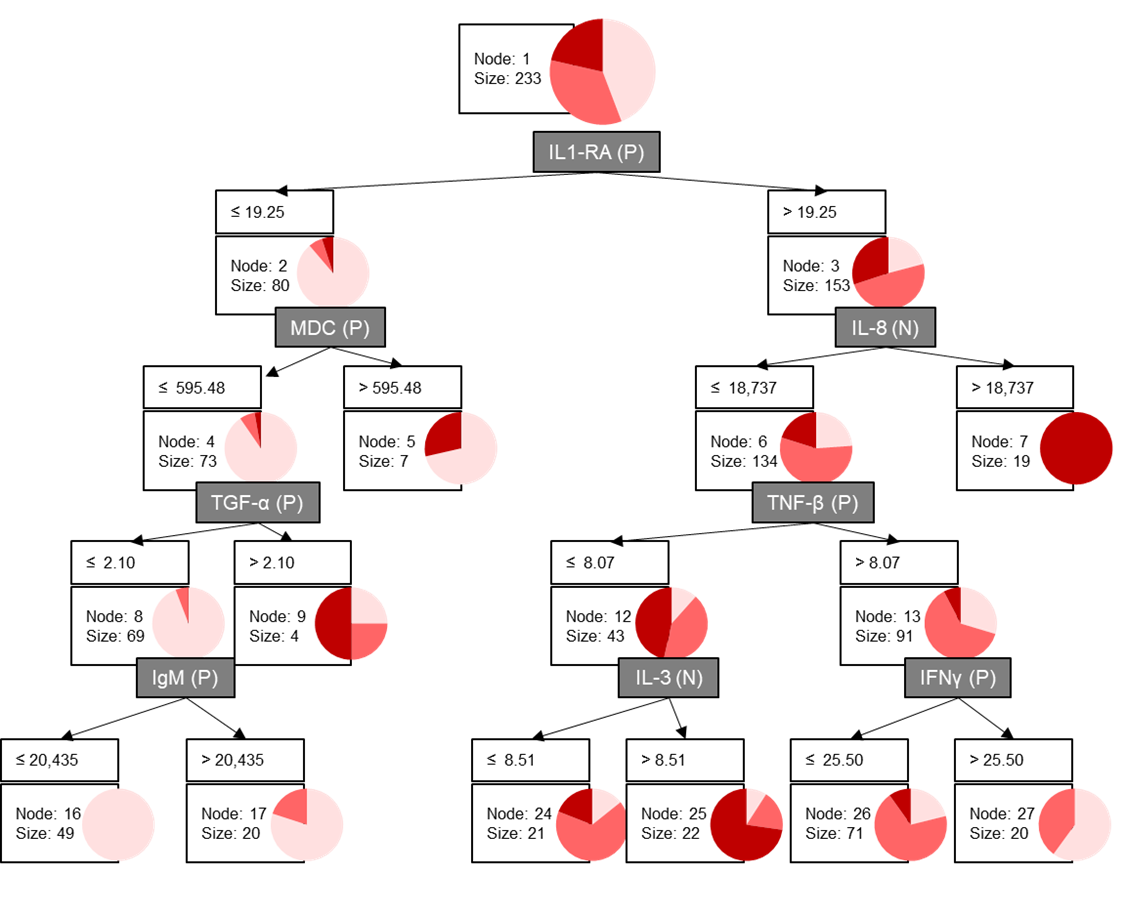

B

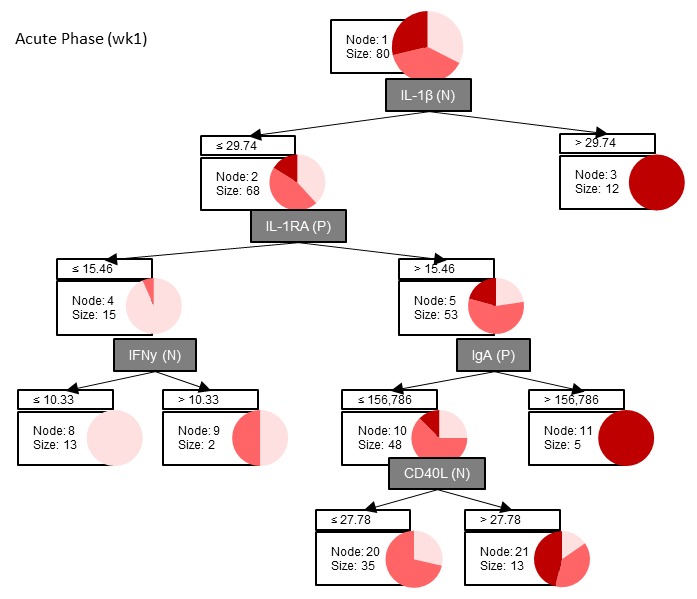

**Figure E9.** Immune predictors for mild, moderate, or severe COVID-19.
